# AP-1B facilitates endocytosis during cell migration

**DOI:** 10.1101/818856

**Authors:** Margaret Johnson Kell, Su Fen Ang, Lucy Pigati, Abby Halpern, Heike Fölsch

## Abstract

The epithelial cell-specific clathrin adaptor AP-1B has a well-established role in polarized sorting of cargos to the basolateral membrane. Here we demonstrate a novel function for AP-1B during collective cell migration of epithelial sheets. We show that AP-1B colocalized with *β*1 integrin in focal adhesions during cell migration using confocal microscopy and total internal reflection fluorescence (TIRF) microscopy on fixed specimens. Further, AP-1B labeling in cell protrusion was distinct from labeling for the canonical endocytic adaptor complex AP-2. Using stochastic optical reconstruction microscopy (STORM) and live TIRF imaging we identified numerous AP-1B-coated structures at or close to the plasma membrane in cell protrusions. Importantly, immuno-electron microscopy (EM) showed AP-1B in clathrin-coated pits and budding vesicles at the plasma membrane during cell migration. Our data therefore established a novel function for AP-1B in endocytosis. We further show that *β*1 integrin was dependent on AP-1B and its co-adaptor, autosomal recessive hypercholesterolemia protein (ARH), for sorting to the basolateral membrane. Notably, we found that expression of AP-1B (and ARH) slowed epithelial-cell migration, and qRT-PCR analysis of human epithelial-derived cell lines revealed a loss of AP-1B expression in highly metastatic cancer cells indicating that AP-1B-facilitated endocytosis during cell migration might be an anti-cancer mechanism.

## INTRODUCTION

Organ cavities are lined with columnar epithelial cells that organize apical domains luminally, whereas basolateral domains are contacting neighboring cells and the basement membrane. This monolayer architecture has to be maintained throughout life to avoid diseases such as metastatic cancer and polycystic kidney disease (Mellman and Nelson, 2008). To ensure this, epithelial cells continuously sort membrane receptors and adhesion molecules to either surface domain (Weisz and Fölsch, 2016; Weisz and Rodriguez-Boulan, 2009). However, little is known about the involvement or behavior of the polarized sorting machinery during epithelial wound-healing.

Major sorting stations for the polarized delivery of cargos to the plasma membrane are the *trans*-Golgi network (TGN) during biosynthesis and recycling endosomes (REs) for sorting of endocytic cargos and biosynthetic cargos that reach REs from the TGN (Ang et al., 2004; Ang and Fölsch, 2012; Fölsch et al., 2009). Basolateral sorting in REs depends on AP-1B (Fölsch, 2015b). AP-1B is a heterotetrameric clathrin adaptor that shares its two large *γ* and *β*1 and its small *σ*1 subunit with AP-1A (Fölsch et al., 1999). The only difference between both complexes is the incorporation of their respective medium subunits µ1B or µ1A. Unlike µ1A, µ1B is tissue-specific and its expression was found restricted to columnar epithelial cells with the exception of the kidney cell line LLC-PK1 that polarizes but does not express µ1B (Ohno et al., 1999). LLC-PK1 cells are thought to be derived from renal proximal tubules that are naturally devoid of AP-1B expression (Schreiner et al., 2010). Since its discovery, LLC-PK1 cell lines with or without exogenous expression of µ1B to restore AP-1B function have been widely used to analyze polarized protein sorting (Fölsch, 2015a). Like other members of the family of heterotetrameric adaptor complexes such as AP-1A, AP-2, AP-3 and AP-4, AP-1B directly recognizes cargos with YxxØ motifs via µ1B and [D/E]xxxL[L/I] motifs, LL-motifs for short, through a shared surface between *σ*1 and *γ* adaptin (Fölsch, 2015b; Ohno et al., 1999; Weisz and Fölsch, 2016).

Noticeably, although highly homologous, we and others found that AP-1A and AP-1B are functionally distinct. Whereas AP-1A is largely localized at the TGN, AP-1B localizes and functions in REs of fully polarized epithelial cells (Fölsch et al., 2001; Gan et al., 2002; Gravotta et al., 2007). Careful biochemical analysis showed that both complexes do not form mixed vesicle populations (Fölsch et al., 2003), although their localization as judged by immunofluorescence may appear mixed (Guo et al., 2013).

AP-1B worked together with the co-adaptor autosomal recessive hypercholesterolemia (ARH) protein for sorting of an artificial cargo with FxNPxY signal (Kang and Fölsch, 2011). However, it remained to be shown if AP-1B and ARH would also co-operate in the sorting of endogenous cargos such as the focal adhesion molecule *β*1 integrin that contains FxNPxY sorting motifs (Moser et al., 2009). The interaction between ARH and AP-1B was mediated via the large *β*1 adaptin subunit and both AP-1A and AP-1B were pulled down by ARH *in vitro*; however, AP-1A did not co-operate with ARH in cells (Kang and Fölsch, 2011). Most likely, the lipid environment present at the TGN that is enriched for phosphatidylinositol 4-phosphate [PI(4)P, (Wang et al., 2003)] may not be amicable for ARH recruitment in contrast to REs. Indeed, AP-1B expression in epithelial cells led to an accumulation of PI(3,4,5)P_3_ in REs, and in turn we also found that AP-1B localization in REs depended on PI(3,4,5)P_3_ (Fields et al., 2010). This recruitment was mediated by µ1B, because mutations in a specific region of its cytosolic domain (R338N/N339S/V340E) that aligns with the PI(4,5)P_2_ binding patch of µ2 (Collins et al., 2002) led to a loss of RE localization (Fields et al., 2010). In addition to PI(3,4,5)P_3_, we identified Arf6 as important for AP-1B function (Shteyn et al., 2011). Arf6 precipitated AP-1A and AP-1B in *in vitro* pull downs, and mutant Arf6 disrupted AP-1B sorting function in polarized cells (Shteyn et al., 2011). Exogenous over-expression of Arf6 in LLC-PK1 cells grown in cell clusters induced ruffling. Importantly, only AP-1B but not AP-1A was recruited into these membrane ruffles (Shteyn et al., 2011). However, the role that AP-1B might play in membrane ruffles remained elusive.

It is generally accepted that whereas AP-2 facilitates clathrin-mediated endocytosis, all other heterotetrameric AP complexes facilitate sorting at the TGN or in endosomes (Boehm and Bonifacino, 2001; Hirst and Robinson, 1998; Nakatsu and Ohno, 2003). However, studies on ubiquitously-expressed adaptor complexes are typically not performed in epithelial cells. Moreover, studies on AP-1B are typically performed in polarized cells grown in monolayers or cell clusters that are devoid of membrane ruffles. In this study, we demonstrate that in addition to its role in REs, AP-1B localizes to the plasma membrane in cell protrusions to facilitate endocytosis during cell migration in wound healing assays. This was not seen before, because LLC-PK1 cells typically don’t form membrane ruffles when grown in clusters or as monolayers for multiple days before immunofluorescence staining. However, LLC-PK1 cells start to ruffle and migrate when induced through Arf6 expression (Shteyn et al., 2011) or wounding of monolayers (this study). Importantly, AP-1B localized in areas at the plasma membrane that were distinct from AP-2-positive areas. We further found that *β*1 integrin was an AP-1B and ARH cargo protein, and *β*1 integrin colocalized with AP-1B in cell protrusions during cell migration. Moreover, expression of AP-1B and ARH slowed the speed of collective cell migration. Only AP-1B but not ARH expression was lost in highly metastatic cancer cells suggesting that AP-1B’s function during collective cell migration might be physiologically relevant.

## RESULTS

### β1 integrin and AP-1B colocalize in cell protrusions

Focal adhesions are formed when integrin heterodimers, consisting of an *α* and a *β* integrin chain, attach to the extracellular matrix (Horton et al., 2016). They are necessary for forward movement, and integrin heterodimers that contain *β*1 integrin are well known for their role in generating speed during cell migration (Horton et al., 2016; Paul et al., 2015). Because we found that AP-1B was recruited into cell protrusions facilitated by Arf6 (Shteyn et al., 2011), we wondered if AP-1B would colocalize with *β*1 integrin during collective migration of epithelial sheets. To aid us in our analysis, we created stable LLC-PK1 cell lines that expressed µ1A or µ1B with internal YFP-tags, because there are no suitable antibodies that would distinguish between µ1A/AP-1A and µ1B/AP-1B by immunofluorescence. The YFP-tags were placed between amino acids 230 and 231 as described previously for HA-tagged variants and guided by the crystal structure of µ2 (Fölsch et al., 2001; Owen and Evans, 1998). The newly created cell lines had modest expression levels as compared to endogenous µ1A, and comparable to the expression levels of the HA-tagged variants in LLC-PK1::µ1A-HA and LLC-PK1::µ1B-HA cell lines that were established earlier [Sup. Fig. 1A, (Fölsch et al., 2001)]. Importantly, anti-GFP immunoprecipitations of µ1A-YFP and µ1B-YFP co-precipitated *γ*-adaptin indicating that they were incorporated into AP-1A-YFP and AP-1B-YFP complexes (Sup. Fig. 1B).

To test if AP-1B might colocalize with *β*1 integrin during cell migration, we wounded LLC-PK1::µ1B-YFP cells grown as monolayers on coverglass coated with 1 mg/ml matrigel. Subsequently, cells were allowed to migrate as sheets for up to six hours before fixation and co-staining for *β*1 integrin as well as the actin cytoskeleton and cell nuclei. LLC-PK1::µ1A-YFP cells served as controls. Remarkably, confocal analysis revealed colocalization of AP-1B and *β*1 integrin at the edge of cell protrusions in front of actin arches. To demonstrate this localization, we imaged 3D galleries throughout the specimens. Subsequently, the individual X-Y slices of the gallery were collapsed into one merged plane in a so-called ‘focused gallery projection’ (Fig. 1B). Importantly, AP-1A did not show this prominent colocalization with *β*1 integrin (Fig. 1A) suggesting that AP-1B may have a function in cell migration that may not be shared by AP-1A.

**Figure 1:**
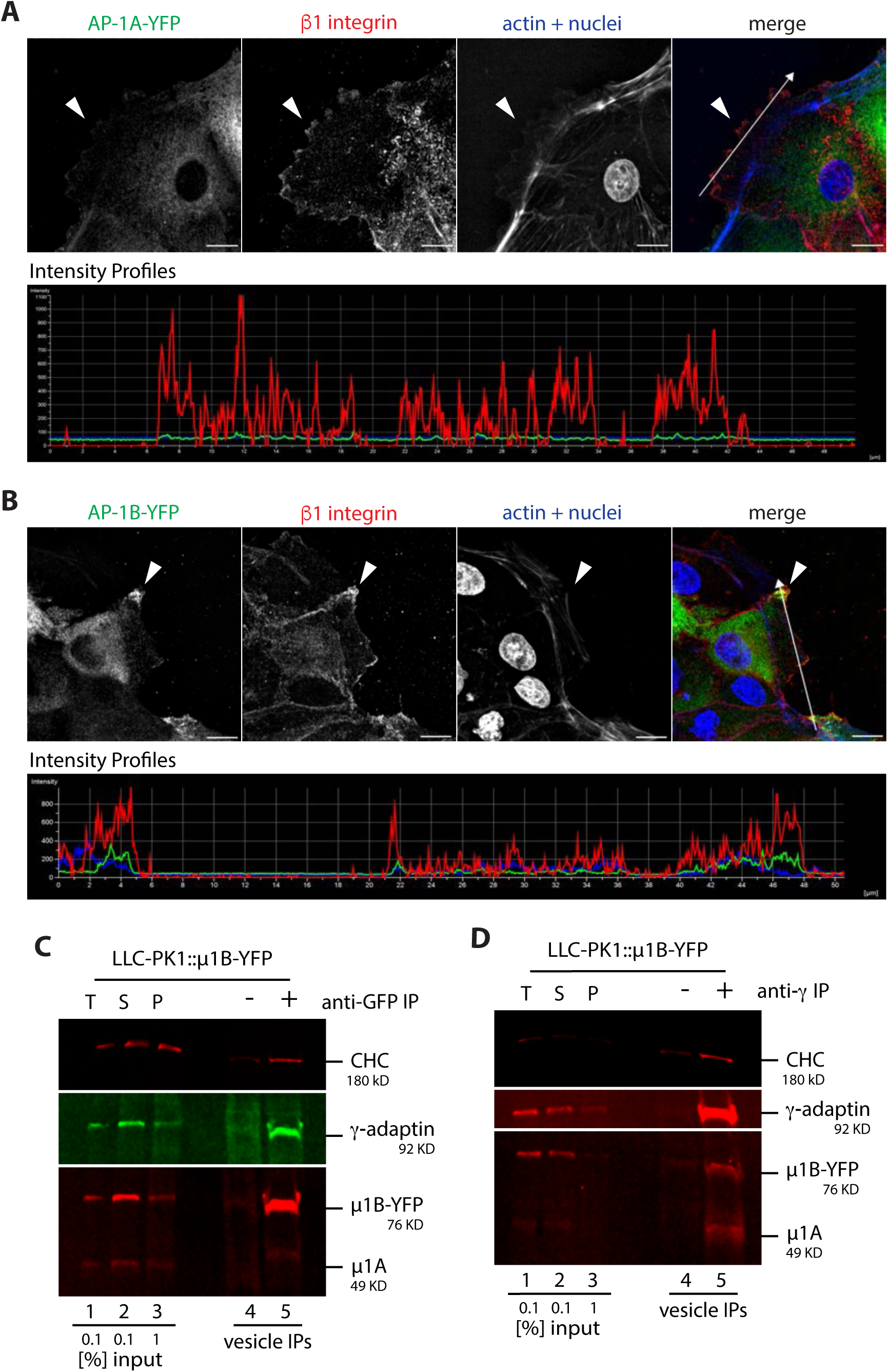
AP-1B is present in cell protrusions. LLC-PK1::µ1A-YFP (A) and LLC-PK1::µ1B-YFP (B) cells were grown on matrigel-coated coverglass for 2 d. Cells were wounded and fixed 6 h later. Specimens were stained for YFP (green), *β*1 integrin (red), actin cytoskeleton (blue), and nuclei (blue), and analyzed by confocal microscopy. Representative collapsed images of acquired 3D galleries are shown in the upper panels. Arrowheads point to the edges of cell protrusions. Arrows in the merged images indicate the area of line scans for intensity profiles shown in the lower panels. Bars are 10 µm. (C) AP-1B-YFP-coated vesicles were immunoisolated out of LLC-PK1::µ1B-YFP cells from crude clathrin-coated vesicle pellets using goat-anti-GFP antibodies (lane 5) or generic goat IgGs as control (lane 4) as described in Material and methods. Samples were analyzed by SDS PAGE and Western blot using anti-CHC and mouse-anti-*γ*-adaptin antibodies as well as antibodies that detect both µ1A and µ1B (anti-µ1A/B antibody). T = 0.1% of the total lysate (lane 1), S = 0.1% of the high-speed (100,000 × *g*) supernatant (lane 2), P = 1% of the high-speed pellet used for the immunoprecipitations (lane 3). (D) AP-1A and AP-1B-YFP-coated vesicles were immunoisolated out of LLC-PK1::µ1B-YFP cells from crude clathrin-coated vesicle pellets using mouse-anti-*γ*-adaptin antibodies (lane 5) or mouse-anti-LDL receptor (C7) antibodies as control (lane 4) essentially as described in (C). Western blots were carried out with anti-CHC, anti-µ1A/B, and rabbit-anti-*γ*-adaptin antibodies.

To test if AP-1B-YFP would still be incorporated into clathrin-coated vesicles during cell migration, we seeded LLC-PK1::µ1B-YFP cells in 20-cm dishes. Monolayers were wounded with multiple scratches to create several wound edges throughout the surface of the plates and allowed to migrate for several hours before cells were homogenized and crude clathrin-coated vesicles were harvested by centrifugation in 1% Triton-X100 (Pearse, 1982). Pellets were resuspended in a sucrose buffer and subjected to immunoisolations (Fölsch et al., 2003) with anti-GFP or anti-*γ*-adaptin antibodies followed by SDS PAGE and Western blot (Fig. 1C & 1D). AP-1B-YFP was brought down in both immunoprecipitations. As expected, anti-GFP antibodies co-precipitated *γ*-adaptin and clathrin heavy chain (CHC, Fig. 1C). Furthermore, anti-*γ*-adaptin antibodies co-precipitated µ1B-YFP and endogenous µ1A as well as CHC (Fig. 1D). We conclude that AP-1B-YFP was incorporated into clathrin-coated vesicles in migrating epithelial sheets. Unfortunately, although we tried multiple different anti-*β*1 integrin antibodies, we were unable to identify one that was sensitive enough to detect *β*1 integrin by Western blot in the cell lysates used for vesicle immunoisolations. Therefore, we could not directly probe biochemically if *β*1 integrin was incorporated into AP-1B vesicles. Regardless, we made the astounding and unexpected discovery that *β*1 integrin colocalized with AP-1B at the plasma membrane in migrating cells.

Next we sought to further investigate AP-1B localization in cell protrusions at higher resolution. To this end, we performed total internal reflection fluorescence (TIRF) microscopy on fixed and stained specimens. TIRF microscopy detects signals that are at or in close proximity to the plasma membrane in an evanescent field at the coverglass-specimen interface that is typically less than 100 nm (Mattheyses et al., 2010). LLC-PK1::µ1B-YFP cells as well as LLC-PK1::µ1A-YFP cells as controls were seeded on 1 mg/ml matrigel-coated MatTek dishes. Monolayers were wounded and cells were allowed to migrate for several hours prior to fixation and staining for YFP, *β*1 integrin, and CHC. The actin cytoskeleton was visualized with Alexa 405-labeled Phalloidin. After completion of staining, specimens were maintained in buffer and imaged on the same day as soon as possible. Wound edges were scanned and well-formed protrusions were imaged. Fig. 2A shows a representative protrusion of an LLC-PK1::µ1A-YFP cell. Note how AP-1A-YFP does not extend to the cell edge that is positive for *β*1 integrin. These *β*1 integrin-positive cell edges are marked by arrowheads in selected insets. Overall, there was little overlap between AP-1A-YFP and *β*1 integrin. This was determined by an object-based colocalization analysis that measured the number of *β*1 integrin-positive and CHC-positive objects within AP-1A-YFP-positive areas of well-formed cell protrusions (see Materials and methods for details). On average we measured 26.2±19.7 *β*1 integrin objects within 100 AP-1A-YFP-positive areas (Fig. 2C). In addition, we measured on average 51.3±25.1 CHC-positive objects within 100 AP-1A-YFP-positive areas (Fig. 2D). Only 7.5% of AP-1A-YFP-positive areas contained both *β*1 integrin and CHC-positive objects (Fig. 2E). In contrast, AP-1B-YFP staining was found at the very edge of the cell where AP-1B-YFP colocalized with *β*1 integrin as indicated by arrowheads in selected insets (Fig. 2B). On average there were 62.8±32.5 *β*1 integrin-positive objects within 100 AP-1B-YFP-positive areas in well-formed cell protrusions (Fig. 2C). This increase in colocalization is statistically significant (P = 0.0022) in comparison to the overlap between AP-1A-YFP and *β*1 integrin. Moreover, there were on average 50.6±23.3 CHC-positive objects in 100 AP-1B-YFP-positve areas (Fig. 2D). This is comparable to the overlap between AP-1A-YFP and CHC (P = 0.1). Lastly, 20.4±16% of AP-1B-YFP-positive areas contained both *β*1 integrin and CHC-positive objects (Fig. 2E, P = 0.0189 in comparison to AP-1A-YFP areas that were positive for both *β*1 integrin and CHC). Note, the overlapping staining of both AP-1A and AP-1B with CHC is most likely underestimated, because a fully-formed clathrin coat is expected to interfere with the ability of anti-GFP antibodies to stain the YFP-tagged µ chains.

**Figure 2:**
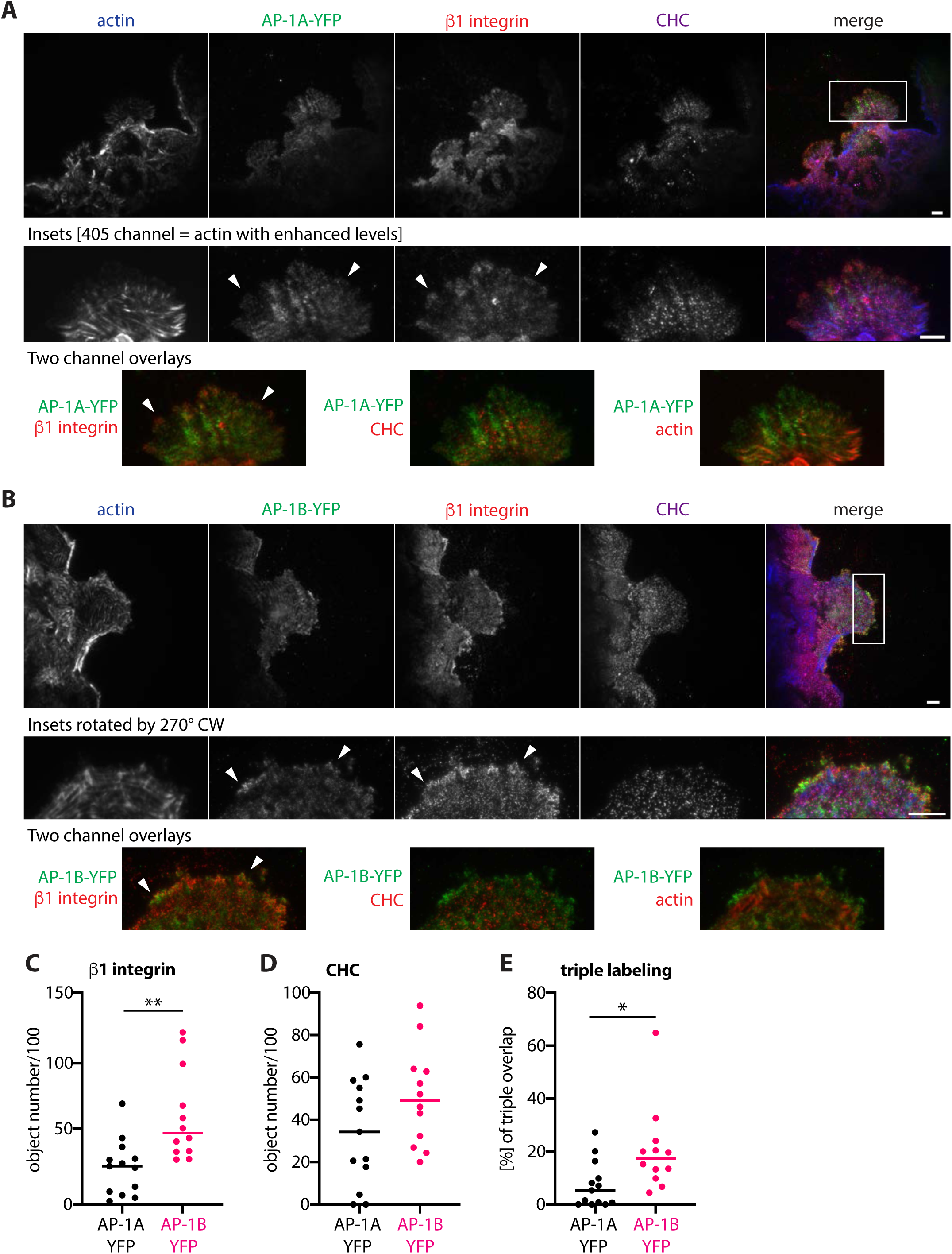
AP-1B colocalizes with β1 integrin in cell protrusions. LLC-PK1::µ1A-YFP (A) and LLC-PK1::µ1B-YFP (B) cells were grown in matrigel-coated MatTek dishes for 2 d, wounded and fixed after cells had well-formed protrusions after 4-6 h. Cells were immunolabeled for YFP (green), *β*1 integrin (red), CHC (magenta), and the actin cytoskeleton (blue). Specimens were imaged by TIRF microscopy and representative images are shown. Rectangles in the merged images indicate areas that were cropped for zoomed-in displays (insets). Arrowheads in the insets point to the *β*1 integrin-positive cell edges. Two-channel overlays were generated in Photoshop. The levels for the 405 channel for the insets in (A) were enhanced in Nikon elements software before exporting the cropped images as TIF files for better appreciation of the actin staining. Bars are 10 µm. (C-E) Object-based colocalization analysis of *β*1 integrin-positive objects (C) and CHC-positive objects (D) in AP-1A-YFP or AP-1B-YFP-positive areas as well as measurement of fields with all three markers (E) was performed with Nikon Elements software as described in Materials and methods. We analyzed 13 well-formed protrusions of LLC-PK1::µ1A-YFP cells from three independent experiments, and 12 well-formed protrusions of LLC-PK1::µ1B-YFP cells from three independent experiments. **, P < 0.003; *, P <0.02.

We conclude that AP-1B-YFP colocalizes with *β*1 integrin in cell protrusions during collective cell migration. This colocalization was not observed for AP-1A-YFP.

### AP-1B does not colocalize with AP-2

AP-2 is well known for its role in facilitating clathrin-mediated endocytosis of integrins (De Franceschi et al., 2016; Moreno-Layseca et al., 2019). Thus we wondered, if AP-1B would colocalize with AP-2. To test this, we stained migrating LLC-PK1::µ1B-YFP cells with antibodies against *α*-adaptin, one of the large subunits of AP-2, as well as the actin cytoskeleton and analyzed the specimens with TIRF microscopy (Fig. 3A). We found virtually no overlap between AP-1B-YFP and AP-2 staining. This is most obvious in the two channel overlay of the insets. Object-based colocalization analysis revealed on average 29.9±25.5 *α*-adaptin/AP-2-positive objects in 100 µ1B-YFP/AP-1B-YFP-positive areas (Fig 3C). Importantly, µ1B-YFP showed colocalization with AP-1’s large *γ*-adaptin subunit (arrowheads in insets, Fig. 3B). On average there were 68.4±27.6 *γ*-adaptin-positive objects within 100 µ1B-YFP-positive areas. The difference in overlap between µ1B-YFP and *α*-adaptin/AP-2 on the one hand and µ1B-YFP and *γ*-adaptin on the other hand was statistically significant (P< 0.0001). Colocalization between µ1B-YFP and *γ*-adaptin is most likely underestimated, because *γ*-adaptin staining labeled both AP-1B-YFP and endogenous AP-1A.

**Figure 3:**
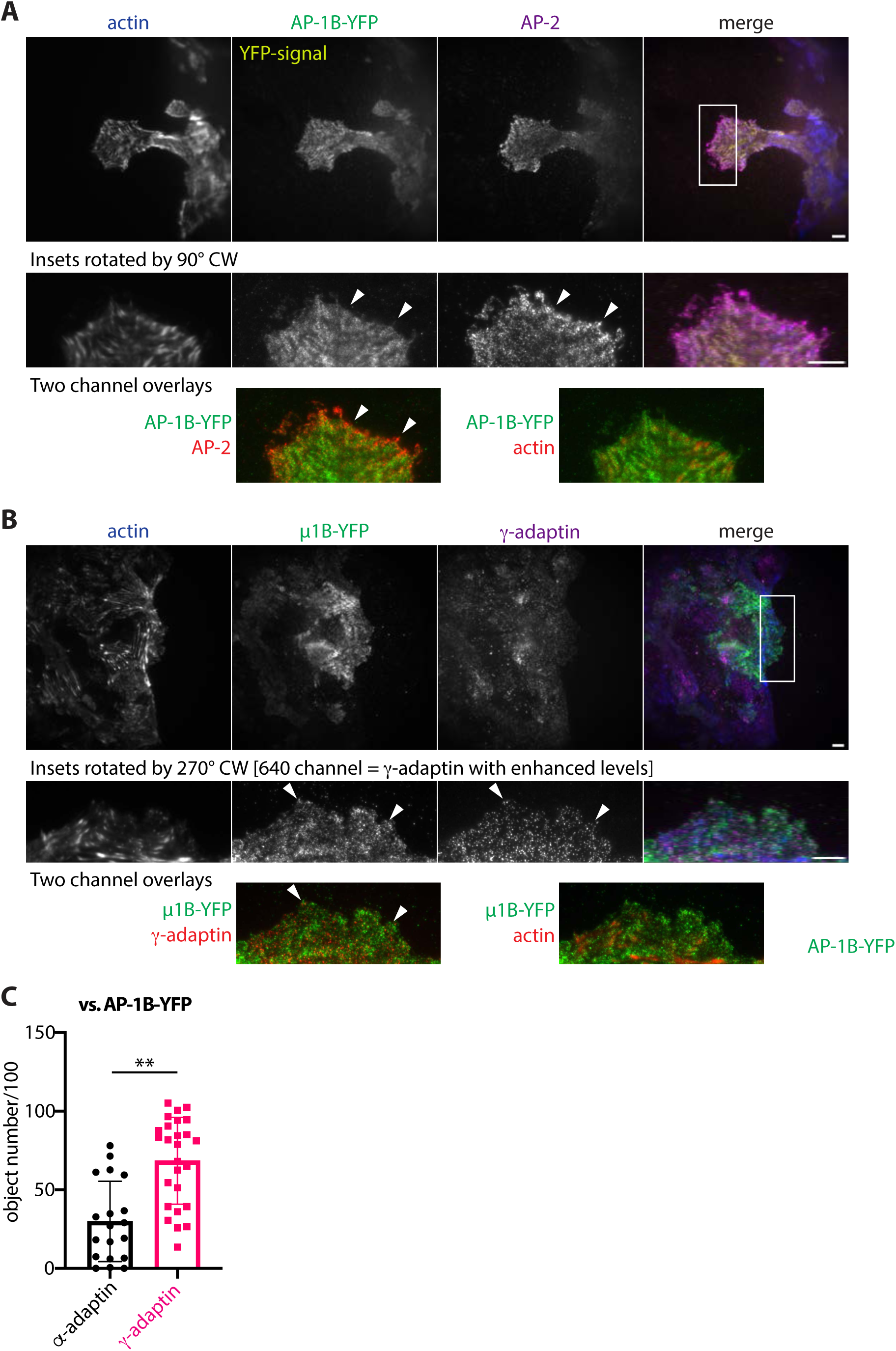
AP-1B-YFP does not colocalize with AP-2. (A) LLC-PK1::µ1B-YFP cells were grown in matrigel-coated MatTek dishes for 2 d, wounded and fixed after cells had well-formed protrusions after 5.5 h. Cells were immunolabeled for AP-2 (magenta) and the actin cytoskeleton (blue). AP-1B-YFP was imaged directly via the YFP signal. Specimens were imaged by TIRF microscopy and representative images are shown. Rectangle in the merged image indicates the area that was cropped for zoomed-in displays (insets). Arrowheads in the insets point to AP-2-positive foci that are devoid of AP-1B-YFP labeling. (B) LLC-PK1::µ1B-YFP cells were grown and treated as described in (A) with fixation 4 h after wounding. Cells were immunolabeled for YFP (green), *γ*-adaptin (magenta), or the actin cytoskeleton (blue). Specimens were imaged by TIRF microscopy and processed as described in (A). Arrowheads in the insets point to µ1B-YFP and *γ*-adaptin-positive foci. The levels for the 640 channel were enhanced in Nikon elements software before exporting the cropped images as TIF files for better appreciation of the *γ*-adaptin staining in the insets. (A/B) Two-channel overlays were generated in Photoshop. Bars are 10 µm. (C) Object-based colocalization analysis of *α*-adaptin/AP-2-positive objects and *γ*-adaptin-positive objects in µ1B-YFP/AP-1B-YFP-positive areas was performed with Nikon Elements software as described in Materials and methods. We analyzed 19 well-formed protrusions of LLC-PK1::µ1B-YFP cells from three independent experiments to score co-localization with *α*-adaptin, and 26 well-formed protrusions of LLC-PK1::µ1B-YFP cells from three independent experiments to score colocalization with *γ*-adaptin. **, P <0.0001.

Next we sought to confirm AP-1B localization in cell protrusions in LLC-PK1::µ1B-HA cells that express HA-tagged µ1B (Fölsch et al., 2001). Cells were again seeded and processed for TIRF analysis on fixed specimens. To this end, specimens were labeled for µ1B-HA/AP-1B-HA, *α*-adaptin/AP-2, CHC, and the actin cytoskeleton. Again, AP-1B-HA was found in cell protrusions in areas that were distinct from AP-2 labeling (Fig. 4, arrowheads “1” pointing to AP-2-positive foci, arrowheads “2” pointing to AP-1B-HA-positive foci). Object-based colocalization analysis revealed on average 15.3±18.4 AP-2-positive objects in 100 AP-1B-HA-positive areas (Fig. 5B). In contrast, the analysis identified significantly more CHC-positive objects (65.8±35.1) in 100 AP-1B-HA-positive fields (P <0.0001, Fig. 4B). In addition, we detected 48.4±30 CHC-positive objects per 100 AP-2-positive fields. This is comparable to the overlap between AP-1B-HA and CHC (P = 0.1557, Fig. 4C)

**Figure 4:**
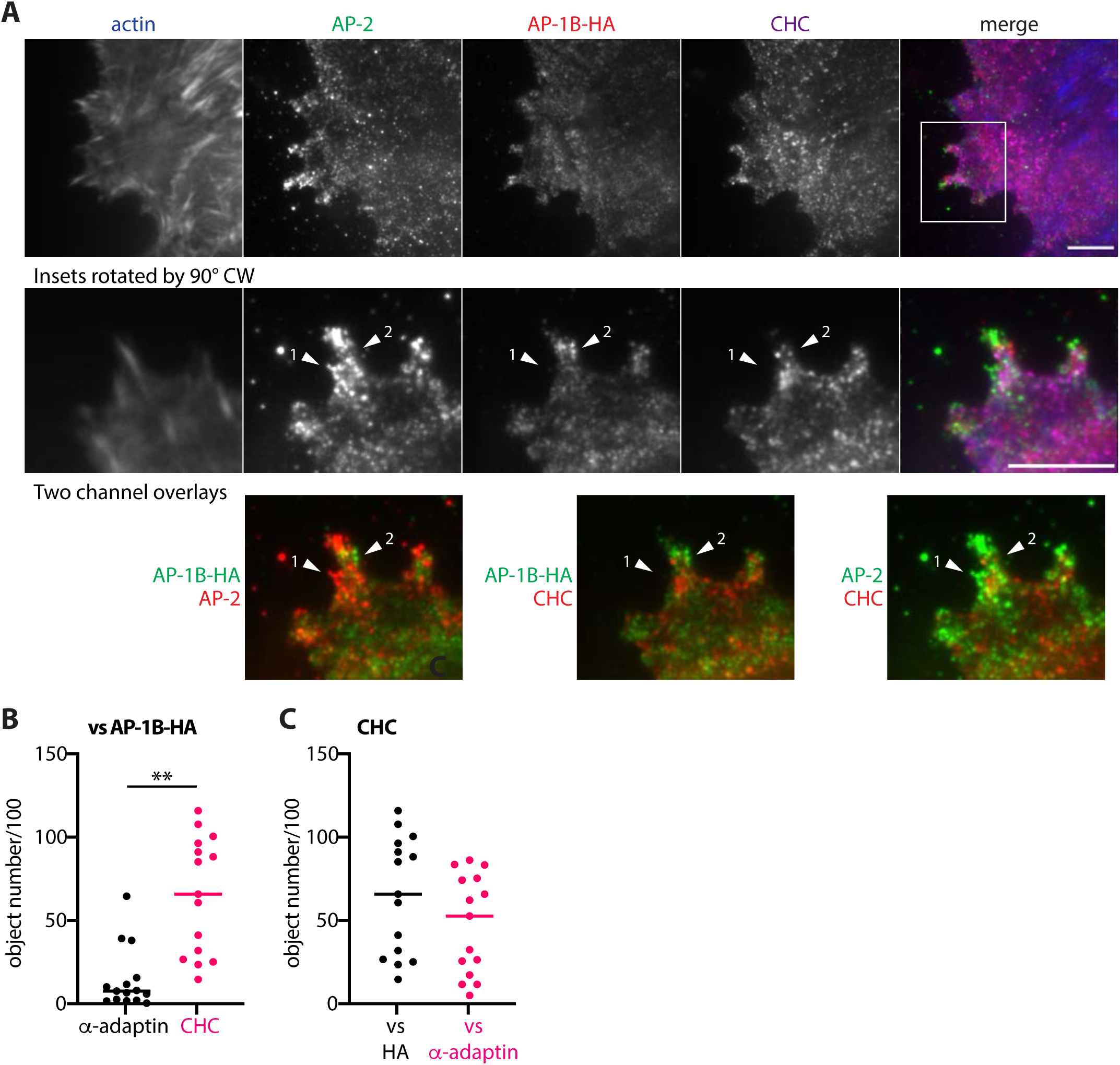
AP-1B-HA does not colocalize with AP-2. (A) LLC-PK1::µ1B-HA cells were grown in matrigel-coated MatTek dishes for 2 d, wounded and fixed after cells had well-formed protrusions after 5.5 h. Cells were immunolabeled for AP-2 (green), µ1B-HA/AP-1B-HA (red), CHC (magenta), and the actin cytoskeleton (blue). Specimens were imaged by TIRF microscopy and representative images are shown. Rectangle in the merged image indicates the area that was cropped for zoomed-in displays (insets). Arrowhead “1” points to AP-2-positive foci and arrowhead “2” points to AP-1B-HA-positive foci. Bars are 10 µm. (B/C) Object-based colocalization analysis of *α*-adaptin/AP-2 and CHC-positive objects (B) in µ1B-HA/AP-1B-HA-positive areas as well as measurement of CHC-positive objects in *α*-adaptin/AP-2-positive areas (plotted against CHC in µ1B-HA/AP-1B-HA-positive areas in [C]) was performed with Nikon Elements software as described in Materials and methods. We analyzed 15 well-formed protrusions of LLC-PK1::µ1B-HA cells from three independent experiments. **, P <0.0001.

**Figure 5:**
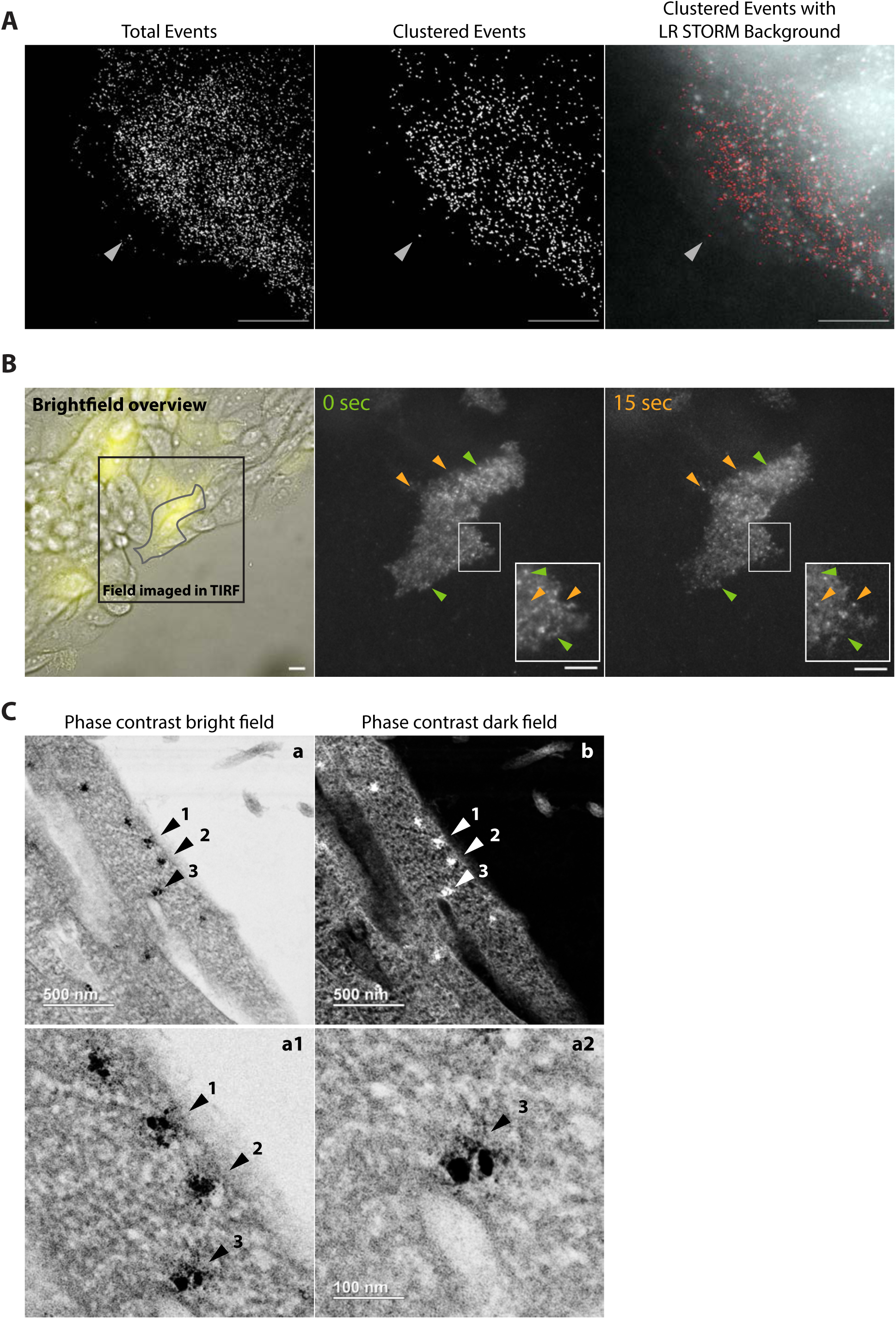
AP-1B localizes in vesicles at the plasma membrane. (A) LLC-PK1::µ1B-HA cells were grown in matrigel-coated MatTek dishes for 2 d. Cells were fixed 1.5 h after wounding and stained with anti-HA antibodies followed by Alexa 647-labeled secondary antibodies. STORM buffer was added and specimens were imaged using STORM followed by processing using Nikon Elements software as described in Materials and methods. We recorded a total of 21 cells in 7 independent experiments. The cell shown had 19,639 out of 21,380 molecules identified. 13,654 molecules were identified in clusters of at least 5 counts in a 70 nm radius. Arrowhead points to a clustered event at the edge of the cell protrusion. LLC-PK1::µ1A cells: no signal in 4 independent experiments. LR, low resolution; bars are 10 µm. (B) LLC-PK1::µ1B-YFP cells were grown in matrigel-coated MatTek dishes for 2 d prior to wounding. Live cell imaging was started 4 h later. The panel on the left shows a brightfield overview with the rectangle showing the area that was imaged using live-TIRF settings (cell area is outlined). Live imaging was carried out for a total of 30 sec with 40 ms exposure times and without delay. Still images at the start (0 sec) and after 15 sec of imaging are shown. Insets show zoomed-in areas that are outlined by rectangles. Green arrowheads mark signals that are present at 0 sec but disappeared by 15 sec, and orange arrowheads mark foci that are present at 15 sec, but not at the start of the movie. Bars are 10 µm. (C) LLC-PK1::µ1B-HA cells were wounded 2 d after seeding in matrigel-coated MatTek dishes. After ∼7 h, cells were fixed and processed for pre-embedding labeling with anti-HA primary and 1.4-nm gold-labeled secondary antibodies, followed by a silver enhancement reaction, and processed for EM as described in Materials and methods. Bright and dark field images were acquired simultaneously with the phase contrast bright field (TE) detector (a) and the high angle annular dark field (HAADF) detector (b) attached to the EM. Zoomed-in images of (a) were either cropped from the TIF file (a1) or directly imaged at higher resolution (a2). Bars are as indicated (500 nm for [a] and [b], 100 nm for [a2]). Because (a1) was cropped out of an existing TIF file it has no scale bar. Numbered arrows point to same regions in the individual images that are positive for AP-1B-HA-labeling in clathrin-coated vesicles (arrowheads ‘1’ and ‘2’) or pits (arrowhead ‘3’) originating at the plasma membrane.

We conclude that AP-1B and AP-2 do not colocalize at the plasma membrane in cell protrusions during epithelial-sheet migration. As expected, both AP-1B and AP-2 colocalized to some extent with CHC.

### Super Resolution Microscopy of AP-1B in cell protrusions

To gather structural information of the AP-1B foci in cell protrusions, we performed stochastic optical reconstruction microscopy (STORM) that relies on detection of stochastic fluorophore ‘blinking’ to obtain high resolution images with a resolution limit of about 20 nm (van de Linde et al., 2011). STORM is carried out in an evanescent field like TIRF microscopy and therefore mainly detects signals originating at or in close opposition to the plasma membrane. For this purpose, LLC-PK1::µ1B-HA cells were grown in monolayers on MatTek dishes coated with 1 mg/ml matrigel. Cells were wounded and allowed to migrate for 1.5 h before being fixed and stained with anti-HA primary antibodies followed by Alexa 647-labeled secondary antibodies. Using STORM, we detected numerous AP-1B-HA-positive events in cell protrusions that had formed at wound edges (Fig. 5A). These signals were scattered indicating AP-1B localization in vesicles as opposed to cytoskeletal structures. Indeed, when events were clustered (5 counts, 70 nm radius to mimic vesicles) many puncta remained (about 70% of events were identified in clusters for the cell shown). This was true for ∼90% of analyzed cells. In contrast, we could not perform this type of analysis with LLC-PK1::µ1A-HA cells, because we virtually detected no STORM signals with this cell line. This suggested that AP-1A was not present at the plasma membrane in this assay. As another internal, negative control we detected no STORM signal in LLC-PK1 cells that were negative for µ1A-HA or µ1B-HA expression.

We conclude that STORM analysis of LLC-PK1::µ1B-HA cells provides further evidence that AP-1B may be present in vesicles located close to or at the plasma membrane during cell migration.

### AP-1B-positive structures are dynamic

If AP-1B indeed would facilitate the formation of endocytic clathrin-coated vesicles, we would expect a highly dynamic behavior of the observed AP-1B structures. Active coating and uncoating of vesicles as well as of vesicles forming and moving out of the evanescent field appear as “blinking”. This is in contrast to labeled endosomes that are tethered to the plasma membrane and thus move laterally in the TIRF field (Cocucci et al., 2012; Lafont et al., 1999). To test if AP-1B engages in vesicle formation at the plasma membrane, we seeded LLC-PK1::µ1B-YFP cells onto MatTek dishes as before. Monolayers were wounded and allowed to recover and migrate for a couple of hours before performing live TIRF imaging. Video S1 shows dynamic appearance and disappearance of AP-1B-YFP signals. The movie was acquired over a total of 30 sec with 40 ms exposure times and imaging without delay. Fig. 5B shows the brightfield overview and still images acquired at 0 sec and 15 sec time points. Green arrowheads point to unique signals at the beginning of the movie and orange arrowheads point to signals at the 15 sec time point indicative of the dynamic changes in labeling patterns. Note that there is virtually no lateral movement of the spots. Instead there is dynamic ‘blinking’. Data analysis that counted ‘spots per frame’ (see Method section for details) revealed that in this particular movie there were 333±60 detected spots per frame. In total we analyzed 20 cells from three independent experiments in this manner and found that dependent on the size of the imaged cell surface area and cell activity there were typically ∼100 to almost 700 spots per frame on average.

Live TIRF analysis thus strengthens our hypothesis that AP-1B-YFP forms clathrin-coated vesicles at the plasma membrane, because the observed dynamic changes in the labeling pattern is indeed indicative of coating and uncoating of vesicles and of vesicles forming and moving out of the evanescent field, as opposed to labeled endosomes tethered to the plasma membrane, because we hardly observed any lateral movement (Cocucci et al., 2012; Lafont et al., 1999). These endocytic AP-1B vesicles would be in addition to endocytic AP-2 vesicles.

### AP-1B localizes in clathrin-coated vesicles and pits at the plasma membrane

Finally, we performed immuno-electron microscopy (immuno-EM) to investigate if AP-1B may truly be recruited to the plasma membrane in clathrin-coated vesicles and pits. We chose LLC-PK1::µ1B-HA cells for these studies, because we previously had reliable results with immuno-gold labeling of AP-1B-HA (Fölsch et al., 2003; Fölsch et al., 2001). To enhance labeling efficiency and reduce steric hindrance of the gold-labeled secondary antibodies, we used nanogold (1.4 nm)-labeled secondary antibodies followed by a silver enhancement reaction during which silver precipitates around the gold particles. The precipitates present as amorphous, high-contrast structures in the EM images.

We readily found numerous AP-1B-HA-labeled structures using either the bright field or dark field detector attached to the EM in clathrin-coated pits and forming vesicles at the plasma membrane in cell protrusions of migrating cells (Fig. 5C). Clathrin-coated structures are identified through their electron-dense appearance. AP-1B-coated clathrin vesicles that are invaginating from the plasma membrane are indicated by arrows ‘1’ and ‘2’, whereas an AP-1B-positive clathrin-coated pit is indicated by an arrow labeled ‘3’ in Fig. 5C. Because exocytic vesicles would no longer be coated with AP-1B and clathrin at the point of fusion with the plasma membrane, these results are a clear indication of AP-1B’s role in endocytosis.

### Basolateral sorting of β1 integrin depends on AP-1B and ARH

Because AP-1B colocalized with *β*1 integrin containing integrin heterodimers (Figs. 1 & 2), we wondered if *β*1 integrin would be a cargo of AP-1B. Recently, de Franceschi and colleagues demonstrated that selective *α* integrin chains have YxxØ motifs that are recognized by AP-2 (De Franceschi et al., 2016). These *α* chains should also be recognizable by AP-1B (Ohno et al., 1999). In addition, *β*1 integrin has two FxNPxY motifs in its cytoplasmic tail [GENPIY and VVNPKY, Fig. 6A (Moser et al., 2009)] that have been reported to interact with endocytic co-adaptors such as Dab2, numb, and ARH. Previously, we showed that ARH also cooperated with AP-1B in recycling endosomes to facilitate basolateral sorting of a model cargo protein with an FxNPxY motif (Kang and Fölsch, 2011). Therefore, we asked if *β*1 integrin would depend on AP-1B for basolateral sorting in epithelia as a means of testing if *β*1 integrin may be a cargo protein that can be recognized by AP-1B. To this end, we tested basolateral sorting of *β*1 integrin in LLC-PK1 cells stably expressing µ1A (LLC-PK1::µ1A) or µ1B (LLC-PK1::µ1B) seeded on filter supports to allow for polarization. A basolaterally-localized transmembrane protein is deemed an AP-1B cargo if it is sorted correctly to the basolateral membrane in the presence of AP-1B, but missorted in its absence (Fölsch, 2015a). Polarized cells were fixed and stained with antibodies directed against *β*1 integrin. Indeed, whereas *β*1 integrin was correctly sorted to the basolateral membrane in LLC-PK1::µ1B cells, *β*1 integrin was missorted in LLC-PK1::µ1A cells (Fig. 6B). Moreover, *β*1 integrin was also correctly sorted in LLC-PK1::µ1B-HA and LLC-PK1::µ1B-YFP cells (Figs. 6C & 6D) indicating that the internal tags did not interfere with the sorting function of AP-1B. In addition, *β*1 integrin was missorted in MDCK cells depleted of µ1B (Gravotta et al., 2007). Therefore, basolateral sorting of *β*1 integrin depends on AP-1B.

**Figure 6:**
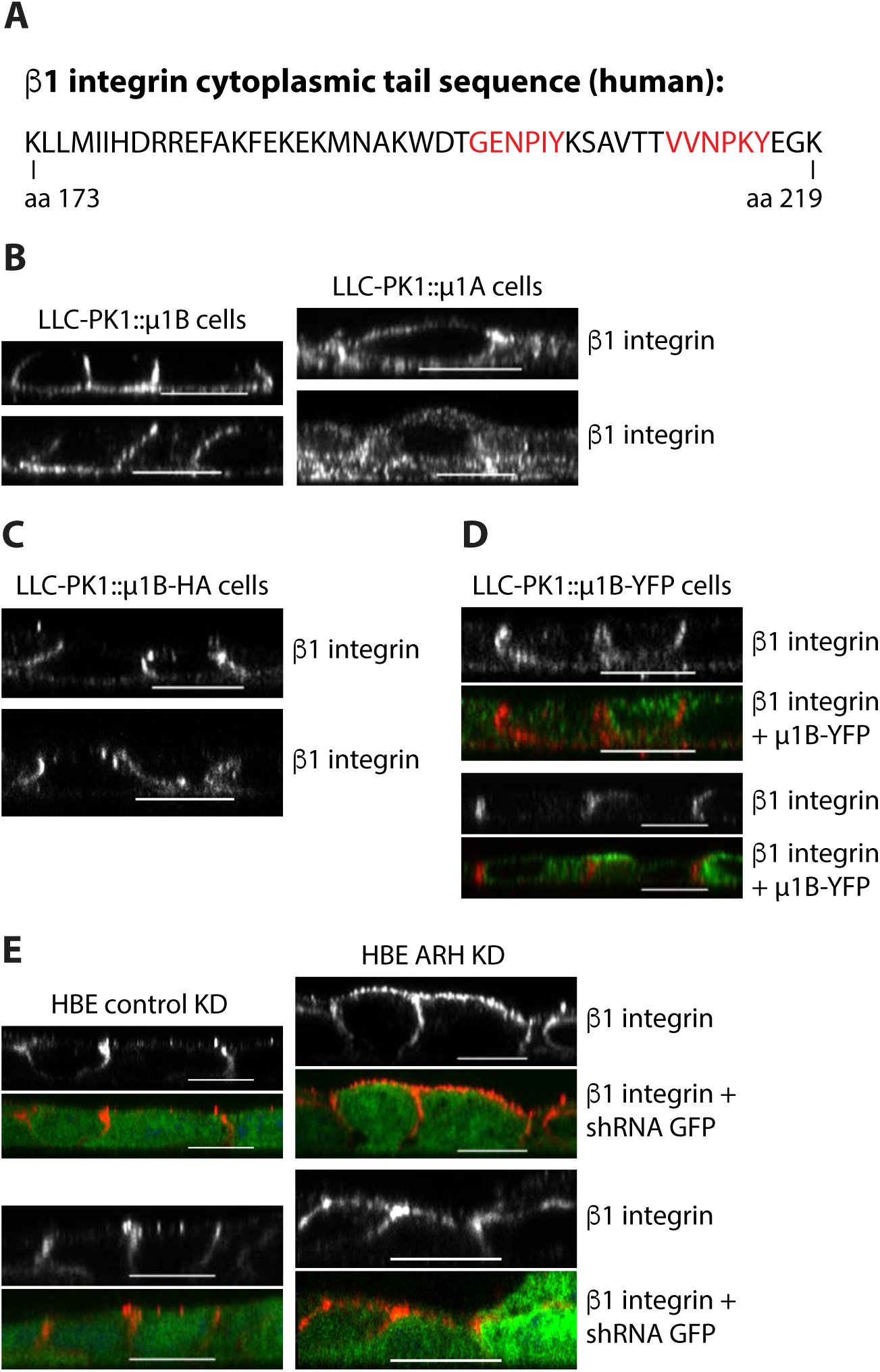
AP-1B and ARH are necessary for β1 integrin sorting. (A) Schematic depiction of the cytoplasmic tail of human β1 integrin. FxNPxY motifs are annotated in red. (B-D) LLC-PK1::µ1B cells (B, left panels) and LLC-PK1::µ1A cells (B, right panels), LLC-PK1::µ1B-HA cells (C), and LLC-PK1::µ1B-YFP cells (D, cells with green signal express µ1B-YFP) were gown on filter supports for 4 d with gentle rocking from side-to-side for 3 d prior to fixation. After fixation, cells were stained for β1 integrin (red in [D]). (E) HBE cells depleted of GAPDH (left panels) or ARH (right panels) were grown on matrigel-coated filter supports for 4 d. Subsequently, cells were fixed and stained for β1 integrin (red). GFP expression indicates shRNA presence. (B-E) Specimens were analyzed by confocal microscopy and representative cross-sections through assembled 3D galleries are shown. Bars are 10 µm.

If the dependence on AP-1B for basolateral sorting of *β*1 integrin is mediated via its FxNPxY motifs than we would predict that ARH should also be necessary for sorting. Previously, we established an ARH knock down protocol in human bronchial epithelial (HBE) cells (Kang and Fölsch, 2011). Thus, to test if ARH plays a role in *β*1 integrin sorting, we depleted ARH or GAPDH as control in HBE cells using shRNA vectors that also express GFP to indicate cells that are positive for shRNA expression. Importantly, whereas *β*1 integrin was correctly sorted in HBE cells depleted of GAPDH, *β*1 integrin was missorted when ARH was depleted (Fig. 6E).

We conclude that basolateral sorting of *β*1 integrin depends on both AP-1B and ARH. Thus, in addition to the previously identified artificial cargo protein LDLR-CT27, we here identify *β*1 integrin as an endogenous cargo for AP-1B and ARH.

### AP-1B expression slows down cell migration

Because *β*1 integrin is important for the generation of speed during cell migration (Paul et al., 2015), and as shown here *β*1 integrin is an AP-1B-dependent cargo protein that colocalized with AP-1B in cell protrusions, we wondered if AP-1B expression would influence the speed of collective cell migration. Thus, we seeded LLC-PK1::µ1A or LLC-PK1::µ1B cells on MatTek dishes that were coated with 1 mg/ml matrigel. Cell monolayers were wounded and migration during wound healing was observed live in a Nikon Biostation for up to 4 hours. Images of selected areas of the wound edge were taken every 15 min (Videos S2 and S3). Fig. 7A shows still images at the beginning, after 2 h and after 4 h of imaging. Using Nikon Elements software, individual cells at the wound edge were tracked manually to determine their traveled path length and distance (Figs. 7B & 7C). On average, LLC-PK1::µ1A cells migrated with a speed of 15.3±2.4 µm/h as opposed to 12.1±1.9 µm/h for cells expressing µ1B, a reduction of 21%. This difference was statistically significant (P <0.002) and was not a result of lost directionality. When migration speeds were determined assessing the ‘straight’ distance, LLC-PK1::µ1A cells traveled at a rate of 12.2±2.8 µm/h whereas LLC-PK1::µ1B cells traveled 8.5±2.4 µm/h (31% reduction in migration speed, P <0.003). Finally, we determined migration prowess by analyzing the total covered area of the migrating cells (Fig. 7D). Whereas LLC-PK1::µ1A cells covered 5.2±1.3% of the total area of the imaged field per h, LLC-PK1::µ1B cells covered only 3.3±1% of the total area per h, a 37% reduction (P <0.002). Although we rarely observed dividing cells during wound healing, growth rates were determined by counting cells at 0, 24, and 48 h after seeding on coverglass coated with 1 mg/ml matrigel. As expected, the growth rates for LLC-PK1::µ1A cells (1.6±0.3) and LLC-PK1::µ1B cells (1.8±0.4) were comparable and therefore not responsible for the observed differences in migration speeds. Furthermore, we detected no differences in the arrangement of actin, microtubule, or keratin cytoskeleton at the leading edge between LLC-PK1::µ1A and LLC-PK1::µ1B cells (not shown, and compare actin staining in Figs. 1 & 2). In conclusion, using three different means of analyzing migration speeds we found that the presence of AP-1B in LLC-PK1 cells slowed down cell migration in wound healing assays.

**Figure 7:**
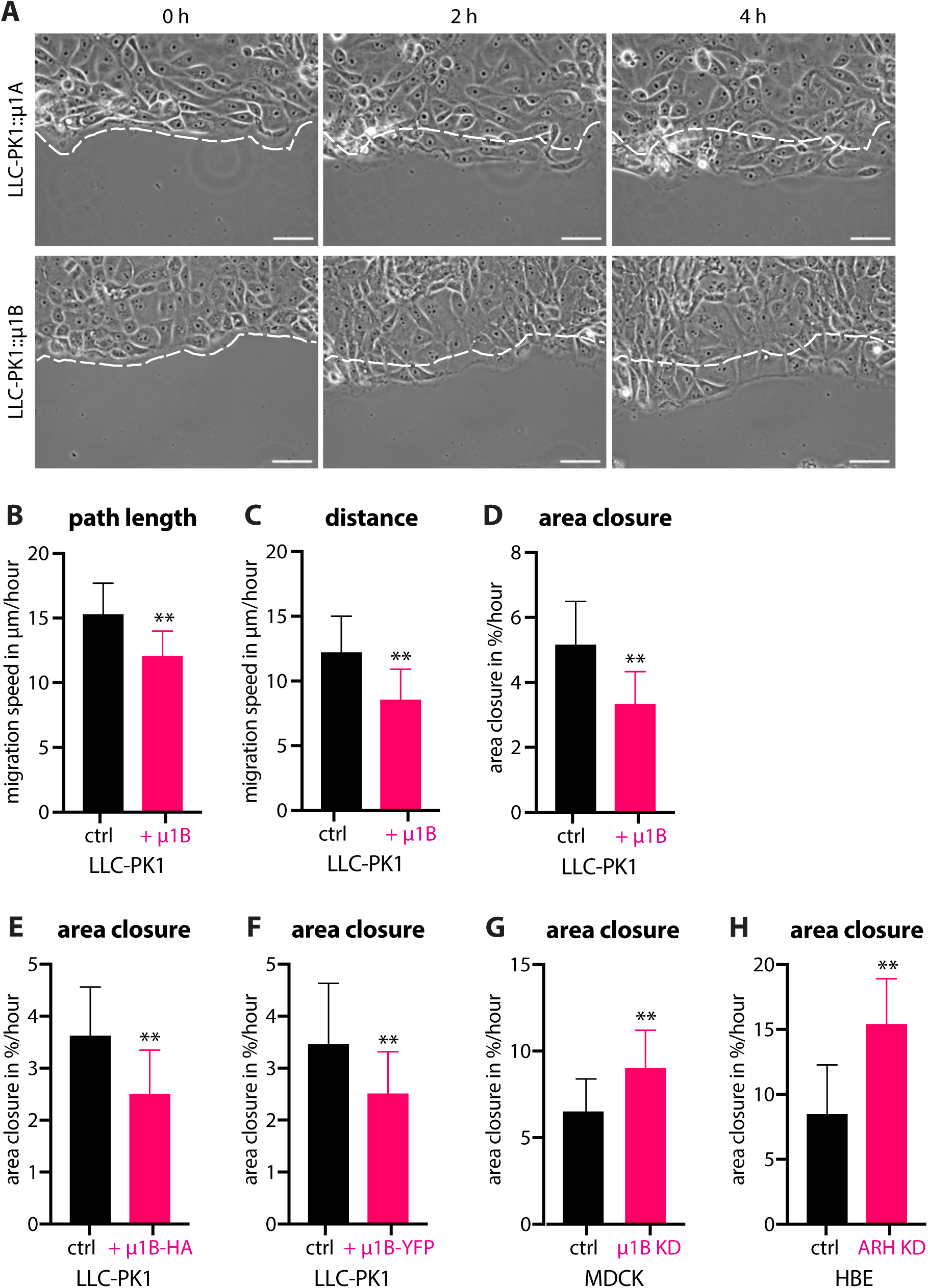
AP-1B expression in epithelial cells slows down migration speeds. LLC-PK1, MDCK, and HBE cells were grown in matrigel-coated MatTek dishes for typically 2 d. After wounding, cells were transferred to a Nikon Biostation for live imaging for up to 4 h. (A) Selected still images of LLC-PK1::µ1A (upper panels) and LLC-PK1::µ1B (lower panels) wounded monolayers at the beginning (0 h), 2 h, and 4 h after the start of data acquisition. Images of selected areas at the wound edge were taken every 15 min. Pixeled lines indicate wound edges at the beginning of live imaging. Bars are 50 µm. (B/C) Traveled path length (B) and distance (C) of migrating LLC-PK1 cells were determined using manual tracking of individual cells at the wound edge as described in Materials and methods. Data represents mean values from three independent experiments. LLC-PK1::µ1A: cells from 9 different areas (a total of 55 individual cells) were analyzed; LLC-PK1::µ1B: cells from 14 different areas (a total of 70 individual cells) were analyzed. (D) Area closure data represents mean values from three independent experiments. LLC-PK1::µ1A: 7 independent areas were analyzed; LLC-PK1::µ1B: 15 independent areas were analyzed. (E/F) Area closure analysis was performed on migrating LLC-PK1 cells stably expressing µ1A-HA or µ1B-HA (E), and µ1A-YFP or µ1B-YFP (F). Data represents mean values from at least three independent experiments. LLC-PK1::µ1A-HA: 15 independent areas; LLC-PK1::µ1B-HA: 32 independent areas; LLC-PK1::µ1A-YFP: 16 independent areas; and LLC-PK1::µ1B-YFP: 22 independent areas were analyzed. (G) Area closure analysis of mock or µ1B-depleted MDCK cells. Data represents mean values from at least three independent experiments using cells from three independent KDs. Control MDCK cells: 23 independent areas; µ1B-depleted MDCK cells: 25 independent areas were analyzed. (H) Migrating HBE cells depleted of ARH or GAPDH as control were analyzed for area closure during wound healing. Area closure data represents mean values of at least three independent experiments using cells from three individual KDs. GAPDH KD HBE cells: 40 independent areas; ARH KD HBE cells: 32 independent areas were analyzed. (B-H) Error bars indicate STDEV. **, P <0.003 with the exception for the comparison of LLC-PK1::µ1B-YFP cells versus LLC-PK1::µ1A-YFP cells for which P was 0.0058. KD = knockdown.

Importantly, the cell lines expressing HA or YFP-tagged µ1B/AP-1B also migrated at a slower pace as compared to the cell lines expressing the correspondingly tagged µ1A/AP-1A. Whereas LLC-PK1::µ1A-HA cells covered 3.6±0.9% of the total area per hour, LLC-PK1::µ1B-HA cells covered only 2.5±0.8% of the total area per hour, a 31% reduction (P=0.0002). Moreover, LLC-PK1::µ1A-YFP cells covered 3.5±1.2% of the observed area per hour compared to 2.5±0.8% coverage achieved by LLC-PK1::µ1B-YFP cells, a 29% reduction (P=0.0058). We conclude that the internal tags did not interfere with AP-1B’s role in cell migration.

To make sure that the effect of AP-1B on cell migration was not specific to LLC-PK1 cells, we depleted µ1B expression in MDCK cells using shRNA using previously established protocols (Anderson et al., 2005; Fields et al., 2010) and determined migration speeds through area closure analysis (Fig. 7G). Whereas mock-depleted MDCK cells covered 6.5±1.9% of the total area per h, µ1B depleted MDCK cells were able to cover 9.0±2.2% of the total area per h, an increase by 38% (P < 0.0001). Thus, using gain-of-function and loss-of-function approaches we found that AP-1B presence in epithelial cells slowed their migration speeds.

Because we found that β1 integrin depended on both AP-1B and ARH for basolateral sorting (Fig. 6), we next asked if ARH also played a role in collective cell migration. Therefore, we seeded HBE cells depleted of ARH or GAPDH as control on MatTek dishes coated with 1 mg/ml matrigel and determined migration speeds in wound healing assays. Area closure analysis showed that whereas control HBE cells only covered 8.5±3.8% of the total area per h, HBE cells depleted of ARH covered almost twice as much area with a speed of 15.4±3.5% of the total area per h, an increase by 81% (P <0.0001, Fig. 7H). This increase in migration speed mimics the increase we saw after µ1B depletion in MDCK cells.

In sum, both the expression of AP-1B and of ARH in AP-1B-positive epithelial cells slowed the speed of migration in wound healing assays.

### AP-1B expression is lost in highly metastatic cancer cells

When we first described AP-1B, we noticed that re-expression of µ1B in LLC-PK1 cells had a profound effect on monolayer appearance. Whereas LLC-PK1::µ1B cell grew in monolayers, LLC-PK1::µ1A cells grew on top of each other (Fölsch et al., 1999). Curiously, *β*1 integrin, unlike other integrins, is localized not only to the basal but also to the lateral surface. Further, it was shown that *β*1 integrin facilitates cell-cell adhesion between keratinocytes and is needed for the invasive behavior of squamous cell carcinomas (Brockbank et al., 2005; Larjava et al., 1990). Therefore, AP-1B expression in epithelial cells may promote tissue homeostasis and may help to prevent cancer metastasis. If this were true, one would expect reduced levels of µ1B/AP-1B in cancer cells. To test this idea, we measured µ1B transcript levels by qRT-PCR in various human cell lines. Indeed, we found µ1B transcripts in normal breast epithelial cells MCF10A and 76NTER, as well as HBE and Caco-2 cells. However, we detected virtually no µ1B transcripts in MDA-MB-231 and HeLa cell lines that were derived from highly metastatic cancers (Fig. 8A). In contrast, ARH transcripts were found in all tested cell lines (Fig. 8B). We conclude that AP-1B may indeed have a positive function in the homeostasis of epithelial tissues.

**Figure 8:**
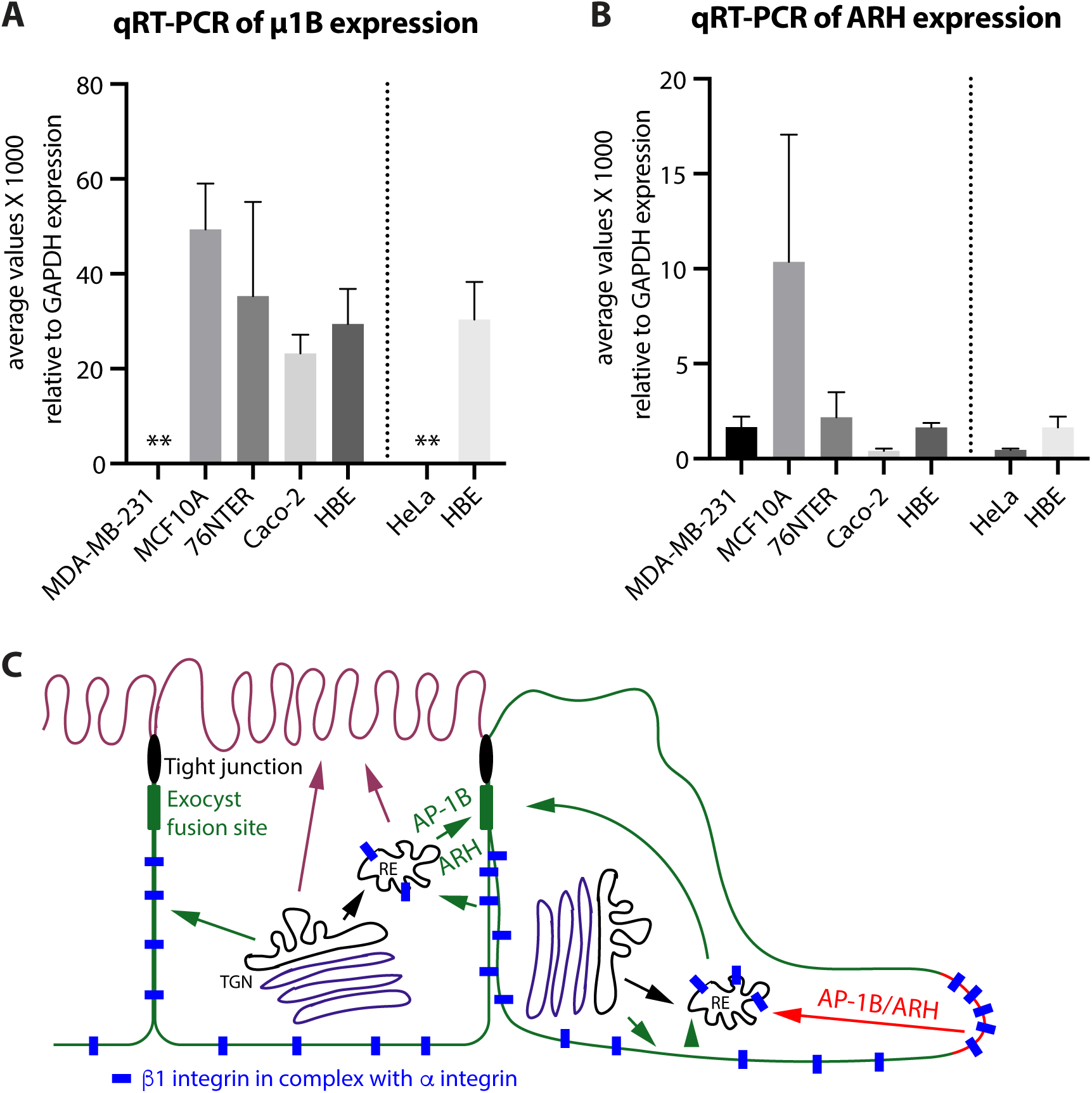
AP-1B expression is lost in highly metastatic cancer cells. (A/B) qRT-PCR was performed using one set of primers specific for human µ1B (A) or human ARH (B). Transcript levels from three independent experiments done in triplicates were plotted in relation to GAPDH transcript levels. Error bars indicate STDEV. **, P <001 (with exception of the pair 76NTER and MDA-MB-231 where P = 0.0372, which is still considered statistically significant). (C) Model depicting the role of AP-1B together with ARH in endocytosis during migration of epithelial monolayers.

### Discussion

In conclusion, we provide data that showed AP-1B colocalizing with *β*1 integrin and clathrin at the plasma membrane in cell protrusions in areas that were not occupied by AP-2. These areas seemed to be composed of vesicles as indicated by STORM and live TIRF microscopy. Further, EM analysis showed AP-1B localizing in clathrin-coated pits and vesicles forming at the plasma membrane in migrating cells establishing AP-1B as an endocytic adaptor. Wound healing assays revealed that both AP-1B and ARH expression in epithelial cells slowed collective cell migration. Further, both proteins were necessary for the basolateral delivery of *β*1 integrin establishing *β*1 integrin as an AP-1B and ARH cargo protein. Taken together, these data suggests that AP-1B may facilitate endocytosis of *β*1 integrin – and most likely other cargo proteins – in cell protrusions thereby removing this cell-matrix adhesion molecule from the leading edge and thus slowing down cell migration (compare Fig. 8C). This may or may not be in addition to a putative role of AP-1B in cargo delivery to focal adhesions or the plasma membrane in general during cell migration.

The fold change in migration speeds in our wound healing assays typically ranged from about 20 to 50%. ARH depletion in HBE cells resulted in a greater change, most likely because ARH affects trafficking of fewer cargos as opposed to AP-1B (Kang and Fölsch, 2011). E-cadherin is one such cargo that depends on LL-signals for basolateral sorting, and sorting efficiency depends on AP-1B (Ling et al., 2007). Notably, the speed of epithelial-sheet migration in wound healing assays is positively correlated with E-cadherin expression at the basolateral membrane (Hwang et al., 2012). Thus, based on these data one would have expected that AP-1B-expressing epithelial cells would migrate at greater speeds as opposed to the corresponding cells without AP-1B expression. This would be the exact opposite of what we found and might explain why the fold-change we observed in migration speed was much greater after depletion of ARH in HBE cells as opposed to depletion of µ1B in MDCK cells. Overall the fold-changes observed are in line with other migration studies done in epithelial cells. For example, depletion of E-cadherin in Caco-2 cells reduced migration speeds by approximately 50% (Hwang et al., 2012), and cleavage of the v-SNARE Vamp3 in MDCK cells stably expressing tetanus neurotoxin resulted in a reduction in velocity from about 17 µm/h to 8 µm/h (Proux-Gillardeaux et al., 2005). Vamp3 is needed for endosomal membrane trafficking, integrity of REs, and AP-1B-facilitated exocytosis in epithelial cells (Fields et al., 2007).

AP-1B may aid in preventing abnormal growth by restricting *β*1 integrin at the basolateral membrane at steady-state and by helping to internalize *β*1 integrin from the plasma membrane in cell protrusion during wound healing. Apical delivery of *β*1 integrin in the absence of AP-1B may be facilitated by galectin-3 (Hönig et al., 2018), and apically expressed *β*1 integrin could potentially help the survival of cells that are extruded apically from epithelial monolayers by promoting cell-cell adhesion at the apical membrane at steady-state (Larjava et al., 1990). During cell migration, high levels of *β*1 integrin at the leading edge may facilitate intercalation of cells when wound edges close potentially leading to abnormal growth in the absence of AP-1B (Brockbank et al., 2005). The fact that histological studies suggested that >80% of kidney cancers arose from proximal tubules that lack AP-1B expression supports this hypothesis (Gu et al., 1991; Holthofer et al., 1983; Martensson et al., 1995; Schreiner et al., 2010). Future investigations will be necessary to determine how AP-1B’s role during cell migration is regulated between its ‘regular’ function in exocytosis from REs and its novel function in endocytosis.

### Putative regulation of AP-1B’s endocytic activity

The leading edges of migrating cells are enriched in PI(3,4,5)P_3_ (Franca-Koh et al., 2007). Furthermore, PI(4,5)P_2_ is generated at focal adhesions and membrane ruffles (Ling et al., 2006). Therefore, leading edges provide favorable environments for the recruitment of both AP-2 and AP-1B. However, we found that AP-1B and AP-2 did not colocalize. Indeed, membrane recruitment of AP-2 is strictly dependent on PI(4,5)P_2_ and thus AP-2 should be absent from areas that are enriched in PI(3,4,5)P_3_ (Collins et al., 2002). In contrast, AP-1B is recruited by PI(3,4,5)P_3_ (Fields et al., 2010), most likely in addition to Arf6 (Fölsch, 2015b; Shteyn et al., 2011) thus explaining why AP-2 and AP-1B localize to different areas in cell protrusions. In contrast, AP-1A is recruited onto PI(4)P-positive membranes and therefore absent from focal adhesions and the plasma membrane (Wang et al., 2003). In order for AP-1B to function in endocytosis, we predict that in addition to PI(3,4,5)P_3_, active Arf6 may also play a role in recruiting AP-1B to the plasma membrane (Santy and Casanova, 2001; Shteyn et al., 2011). Currently, we are unable to easily and reliably co-stain for AP-1B and PI(3,4,5)P_3_ and Arf6, as well as ARH during cell migration due to low transfection efficiencies. To this end, the establishment of new stable cell lines would be required, which is beyond the scope of this study. Regardless of these limitations, our study clearly demonstrates that AP-1B is recruited to the plasma membrane of migrating cells.

Recruitment of AP-1B to the plasma membrane does as far as to our knowledge not happen during steady-state. Even though the basolateral membrane in MDCK cells is enriched in PI(3,4,5)P_3_ (Gassama-Diagne et al., 2006), AP-1B does not localize there (compare µ1B-YFP/AP-1B-YFP fluorescence in Fig. 6D). Previously, we found that the PH-domain of Grp1 that bound both PI(3,4,5)P_3_ and activated Arf6 (Di Paolo and De Camilli, 2006; DiNitto et al., 2007) localized in REs and membrane ruffles, but not at the lateral membranes (Fields et al., 2010). This indicates that perhaps Arf6 may not be activated at cell-cell borders and therefore AP-1B may not be recruited. Indeed, whereas Arf6 was shown to function in endocytosis from the apical membrane (Altschuler et al., 1999), it played no role in endocytosis from the basolateral domain of fully polarized MDCK cells (Boulant et al., 2011).

### Components of the clathrin machinery in focal adhesions

Recent developments in the field emphasized the existence of adhesion complexes that lack association with the cytoskeleton as well as with integrin-associated adhesion components. Indeed, reticular adhesions were described for *α*V*β*5 attachment to substratum in ‘plaques’ that were often much less enriched in actin but were enriched in AP-2 and CHC (Lock et al., 2019). These plaques were hypothesized to prevent clathrin-coated vesicle formation by preventing membrane bending (Baschieri et al., 2018). These types of adhesions could potentially be formed by any integrin in contact with the extracellular matrix (Lock et al., 2019). Moreover, AP-2 regulated adhesion in a 3D matrix independently from CHC by forming tube-like lattices around collagen fibers (Elkhatib et al., 2017). These interactions were most likely driven by recognition of a selected *α* integrin subunit to AP-2 through their YxxØ motif (De Franceschi et al., 2016). Because such YxxØ signals would also be recognized by AP-1B, AP-1B may interact with integrin heterodimers in more than one way during cell migration. Notably, components that facilitate clathrin-mediated endocytosis were found in the meta-adhesome, but not the consensus adhesome, of focal adhesions in addition to Arf GEFs and GAPs and AP-1 subunits (Horton et al., 2015; Lock et al., 2019). In addition, lipid kinases such as PIPKI*γ* were also associated with the adhesome (Winograde-Katz et al., 2014). This strengthens our hypothesis that AP-1B may have a function in focal adhesion biology, because it is predicted that Arfs play a role in AP-1B recruitment (Shteyn et al., 2011), and it has been shown that AP-1B forms tertiary complexes with PIPKI*γ*-90 and E-cadherin (Ling et al., 2007).

Focal complexes are nascent adhesions that are mainly found at the very end of cell protrusions that are not yet engaged with actin, but may develop into mature focal adhesions over time (Moreno-Layseca et al., 2019; Winograd-Katz et al., 2014). Focal adhesion are marked by their attachment to actin stress fibers. They are associated with the integrin adhesome and are generating traction forces during cell migration (Moreno-Layseca et al., 2019). The adhesome contains proteins such as talin and kindlins that bind to the NPxY motifs in *β* integrin tails (Moser et al., 2009), thereby occupying the potential binding sites for ARH. Because of this, and the fact that we found AP-1B localized at the very edge of cell protrusion in a staining pattern that was largely distinct from actin-labeling, we hypothesize that AP-1B may internalize *β*1 integrin-containing heterodimers that are not yet in mature focal adhesions. This would explain why AP-1B-facilitated endocytosis reduces migration speeds. Future studies are planned to analyze if indeed AP-1B associates with nascent adhesions.

## MATERIALS AND METHODS

### Antibodies and Labeling Dyes

Rabbit anti-CHC (ab21679), rabbit anti-GFP (ab290), goat anti-GFP (ab6673), rabbit anti-*α*-adaptin (ab189995), and rabbit anti-*γ*-adaptin (ab220251), as well as CytoPainter phalloidin-iFlour 405 (ab176752) were purchased from Abcam. Mouse anti-*β*1 integrin (MEM-101A) was purchased from Novus Biologicals, mouse anti-HA (16B12) from Covance, and mouse anti-*γ*-adaptin (100/3) from Sigma. Hybridomas producing antibodies against LDL receptor (C7) were purchased from the American Type Culture Collection. Unlabeled, generic goat anti-rabbit IgG antibodies were purchased from Zymed. A polyclonal antibody cross-reacting between µ1A and µ1B was a kind gift from Dr. Linton Traub (University of Pittsburgh, Pittsburgh, PA).

Secondary antibodies labeled with Alexa dyes donkey anti-mouse Alexa 647, donkey anti-mouse Alexa 568, goat anti-mouse Alexa 488, donkey anti-goat Alexa 680, and donkey anti-goat Alexa 488 were purchased from Molecular Probes/Thermo Fisher Scientific. Donkey anti-mouse IRDye 800CW and donkey anti-rabbit IRDye 680RD antibodies were purchased from Li-Cor. Donkey anti-rabbit Cy5 antibodies, peroxidase conjugated goat anti-mouse and goat anti-rabbit antibodies were from Jackson ImmunoResearch. Goat anti-mouse antibodies labeled with 1.4-nm colloidal gold as well as HQ Silver enhancement kit were from Nanoprobes. DAPI solution was purchased from BD Biosciences.

### Cell Culture

All cells were grown at 37°C in the presence of 5% CO_2_. Their respective media were – in general – supplemented with 2 mM L-glutamine, 0.1 mg/ml penicillin/streptomycin, and 0.01 mg/ml ciprofloxacin. LLC-PK1 cells stably expressing µ1A, µ1B, µ1A-HA, µ1B-HA, µ1A-YFP, or µ1B-YFP were maintained in *α*MEM (5% fetal bovine serum) containing 1 mg/ml geneticin. MDCK and HBE cells were grown in MEM (5% fetal bovine serum). HeLa cells were maintained in DMEM (10% fetal bovine serum), MDA-MB-231 were maintained in *α*MEM (10% fetal bovine serum), Caco-2 cells were grown in DMEM (20% fetal bovine serum) supplemented with 0.1 mM non-essential amino acids, 1 mM sodium pyruvate, and 10 µg/ml transferrin. MCF10A and 76NTER cells were maintained in DMEM/F12 medium (5% horse serum) supplemented with 20 ng/ml EGF, 0.5 mg/ml hydrocortisone, 100 ng/ml cholera toxin, 10 µg/ml insulin, and 0.1 mg/ml penicillin/streptomycin. HBE and Caco-2 cells were maintained on coated surfaces. To prepare the coating solution, LHC basal medium was supplemented with 10 mg/ml bovine serum albumin, 3 mg/100 ml bovine collagen I, and 1 mg/100 ml fibronectin.

For wound healing assays, cells were seeded at densities ranging from 9 to 11 × 10^5^ cells onto 22×22-mm coverglass placed in 35-mm dishes (confocal analysis) or into MatTek dishes (35-mm with 14-mm microwell No #1.5 coverglass for Biostation, fixed and live TIRF microscopy, STORM, and EM analysis) typically two days prior to wound healing assays such that cells formed monolayers without being overgrown. Coverglass was acid washed and coated with 1 mg/ml matrigel. Cell monolayers were scratched with a p200 tip to wound them, followed by two washes in growth media. For live-TIRF imaging, cells were grown in media without phenol red. Cells were typically fixed 3-6 h after wounding (1.5 h after wounding for STORM). Live imaging using the Biostation typically started 0.5 to 1 h after wounding, and live-TIRF imaging typically started 4.5 h after wounding.

To monitor polarized sorting of *β*1 integrin, we seeded 4 × 10^5^ cells on 12-mm filter supports (0.4 µm pore size; Corning) and cultured them for 4 days with changes of the medium in the basolateral chamber daily. LLC-PK1 cells were typically put on a rocker platform and rocked slowly and continuously for about 3 days prior to fixation. HBE cells were seeded on filter supports that were coated with 1 mg/ml matrigel.

### Cloning, Retroviruses, and gene knockdown

The internal YFP-tags in µ1A and µ1B were introduced between amino acids 230 and 231 using PCR at the exact same position into which we previously introduced HA-tags (Fölsch et al., 2001). We first amplified N-terminal fragments (amino acids 1 to 230) of µ1A and µ1B with flanking EcoRI at the N-terminus and XhoI and ClaI restriction sites at the C-terminus using the following primer pairs 5’-GCGC GAATTC ATG TCC GCC TCG GCT GTC TTC ATT-3’ and 5’-GCGC ATCGAT CTCGAG TGA TTT GTT CTT GCT GCG GCC AGT-3’ for human µ1B, and 5’-GCGC ATCGAT CTCGAG TGA CTT GCT CTT CCC TCG GCC TGT-3’ and 5’-GCGC GAATTC ATG TCC GCC AGC GCC GTC TAC GTA-3’ for mouse µ1A. We used these fragments to exchange them with the N-terminal parts including the HA-tag of our previous constructs through a simple cut-and-paste approach. We then amplified YFP using pEYFP-C1 as a template and the following primers 5’-GCGC CTCGAG ATGGTGAGCAAGGGCGAGGAG-3’ and 5’-GCGC ATCGAT CTTGTACAGCTCGTCCATGCC-3’. EYFP was introduced between XhoI and ClaI sites in µ1A and µ1B using cut-and-paste technology. The pCB6 vector backbones now containing µ1A-YFP and µ1B-YFP were subsequently used to generate stable cell lines.

The shRNAmir constructs in the lentiviral pGIPZ vector targeting human ARH (CGCTTGGCACTTTAAAGCATTATAGTGAAGCCACAGATGTATAATGCTTTAAAGTGCCAAGC) and human GAPDH were described previously (Kang and Fölsch, 2011; Nokes et al., 2008). The retroviral constructs in RVH1 specifically targeting canine µ1B (CCCCGCAGTCAGTGGCCAATGGTTTCAAGAGAACCATTGGCCACTGACTGCTTTTTGGAAA) and scrambled control were as previously described (Anderson et al., 2005). HBE cells stably depleted of ARH or GAPDH as well as MDCK cells depleted of µ1B or scrambled control were generated by infecting HBE or MDCK cells with respective retroviruses exactly as previously described (Anderson et al., 2005; Pigati et al., 2013). Cells were maintained in growth media with 12 µg/ml puromycin. Depletion of ARH or µ1B mRNAs was monitored using RT-PCR.

### RT-PCR and qRT-PCR

RNA was typically isolated from confluent monolayers of cells seeded in 10-cm plates using the direct-zol RNA miniprep kit from Zymo Research. 1.5 µg of purified RNA was used for reverse transcription using Superscript III enzyme (Invitrogen). Reverse transcribed RNA was subsequently used for conventional RT-PCR or qRT-PCR using Taq polymerase. qRT-PCR was performed using the light Cycler 480 II real-time PCR machine from Roche and SYBR green detection.

(q)RT-PCR primers to amplify human ARH were N-terminal primer 5’-ATCGTGGCTACAGCTAAGGC-3’ and C-terminal primer 5’-GCCTTAGCTGTAGCCACGAT-3’, human µ1B was amplified using N-terminal primer 5’-TCCCTCCCCGACTCCTAAGT-3’ and C-terminal primer 5’-GTGGCCACCAAGTAGAGGTT-3’, and human GAPDH was amplified using N-terminal primer 5’-ACAGTCAGCCGCATCTTCTTT-3’ and C-terminal primer 5’-CAATACGACCAAATCCGTTGACT-3’. Canine ARH was amplified using N-terminal primer 5’-TTGCCTACATTGCACAGAGC-3’ and C-terminal primer 5’-CTTGGACACCTGCCAAAACT-3’, canine µ1B was amplified using N-terminal primer 5’-TGGGGTCAAGTTTGAGATCC-3’ and C-terminal primer 5’-GCCCGGTAACCTCTTTTCTC-3’, and canine GAPDH was amplified using N-terminal primer 5’-GCCAAGAGGGTCATCATCTC-3’ and C-terminal primer 5’-AGGAGGCATTGCTGACAATC-3’.

qRT-PCRs (Fig. 8) were run three times in triplicates on independently isolated RNA samples and analyzed using LightCycler 480 SW 1.5 software. Data was normalized to GAPDH RNA expression and mean values and standard deviations (STDEVs) were determined. Mean values, STDEV, and *n* values were then used to calculate P-values in unpaired student’s *t* tests using GraphPad QuickCalcs (GraphPad Software). Graphs were prepared using GraphPad Prism software.

### Biostation Imaging and Data Analysis

After wounding, cells were transferred to a Nikon Biostation IMQ equipped with a 20x objective (NA 0.8) and a high-sensitivity 1.3-megapixel cooled monochrome camera for brightfield imaging. Data acquisition typically started 30 min to 1 h after placing the specimens into the Biostation, dependent on how fast the system stabilized at 37°C and 5% CO_2_. Data was acquired with Biostation IM software and processed using Nikon Elements software. Manual single cell tracking throughout time frames to determine distance and pathlength traveled by individual cells in a monolayer at the leading edge of cell protrusions was carried out with Nikon Elements AR3.2. For area analysis Biostation nex files were converted to Nikon nd2 files and batch analysis was run using Nikon Elements 4.5. “Field Measurement” measured the biggest inverted area (i.e. the cell-free area) throughout the time-frames (settings: 1. Cell: low pass filter, auto contrast, edge detection; 2. Only_biggist_invert_cell: invert, fill holes, filter on object area). Frames were checked manually and were discarded if they could not be analyzed with this method. Area closure was determined as a [%] of the total area of the frame that became covered by migrating cells divided by the length of imaging (typically 1.5 to 3 h). To determine statistical significance, mean values and STDEVs were determined from the combined data points of at least three independent experiments. Mean values, STDEV, and *n* values were then used to calculate P-values in unpaired student’s *t* tests using GraphPad QuickCalcs (GraphPad Software). Graphs were prepared using GraphPad Prism software. [%] changes were calculated by setting the control cell data at 100%.

### Immunofluorescence Staining and Fluorescent Imaging

Staining was performed essentially as previously described (Cook et al., 2011; Pigati et al., 2013). Briefly, after washing 3 x with PBS^2+^ (phosphate-buffered saline [2.67 mM KCl, 1.47 mM KH_2_PO_4_] plus 0.901 mM Ca^2+^ and 0.493 mM Mg^2+^), cells were fixed with 3% PFA for 15 min (4% PFA for 20 min for fixed TIRF) at room temperature (RT), followed by a 5-min incubation in PBS^2+^ at RT. If dealing with filter supports, filters were cut out from their holders during this 5-min incubation. Specimens were then blocked for 1-h at RT in 10% goat serum (10% fetal bovine serum if a goat primary antibody was used), 2% bovine serum albumin (BSA), and 0.4% saponin (0.2% saponin for cells seeded on filter supports) in PBS^2+^. Primary antibodies were diluted in block solution without serum. Specimens were incubated with primary antibodies for 1-h at RT (or overnight at 4°C for TIRF) followed by 5 washes over 30 min in block solution without serum. Specimens were then incubated with secondary antibodies and phalloidin – if indicated – diluted in block solution without serum for 1-h at RT, followed by 5 washes over 30 min in block solution without serum (or just PBS for fixed TIRF). If needed, DAPI was added to the last wash. Specimens were dipped in H_2_O and mounted in ProLong Gold (Invitrogen). Specimens for fixed TIRF were kept in PBS at 4°C and imaged the same day. Specimens prepared for STORM were kept in PBS^2+^ at 4°C overnight and imaged the next day.

To obtain 3D galleries, specimens were imaged at RT using Nikon A1 or A1R laser scanning microscopes equipped with plan Apo *λ* 100x Oil objectives (NA 1.45) and Nikon A1plus cameras. TIRF was imaged at RT using a Nikon W1 dual cam spinning disk confocal equipped with an Apo TIRF 100x Oil DIC N2 objective (NA 1.49) and a 95B prime Photometrics camera. As well as a Nikon Perfect Focus Ti2 microscope. Live TIRF was imaged at 37°C in a 5% CO_2_ atmosphere using the same instrument. Live imaging was performed with an exposure time of 40 ms, and imaging without delay for up to 1 min. STORM was imaged at RT using a Nikon X1 spinning disk confocal equipped with an Apo TIRF 100x Oil DIC N2 objective (NA 1.49) and an Andor DU-897 X-6974 camera in STORM buffer (50 mM Tris-HCl, pH 8.0, 10 mM NaCl, 0.5 mg/ml glucose oxidase, 40 µg/ml catalase, 10% glucose, 10 mM mercaptoethylamine [MEA]). Data was analyzed using the Nikon Elements software. STORM analysis included identifying real events using software algorithms to identify blinks of medium brightness and eliminating the first 10 to 20 sec of massive photobleaching from the analysis. Identified events were clustered typically at a density of 5 counts in a 70-nm radius. Live TIRF analysis was performed using Nikon Elements software and included equalizing signal intensities in time through a histogram transformation algorithm. To count ‘spots per frame’ we selected a general analysis processing with smooth denoising. Thresholds were set to choose bright spot detection of spots with a diameter of 0.15 µm and an adjusted contrast of 11. Data was processed and combined for presentation using Adobe Photoshop and Adobe Illustrator.

Fixed TIRF data was quantified using the general analysis tool within Nikon Elements 5.2 software for object-based colocalization. To this end, smooth denoising of data and a rolling ball subtraction (1 µm diameter) was chosen, and thresholds set for the channels to be analyzed to reduce non-specific background. Threshold settings were kept constant within data sets. Objects within a chosen channel were subsequently automatically identified with the program. We typically set the channel that detected µ1A or µ1B as the reference channel. For example, if µ1B-YFP was stained with 488-labeled secondary antibodies, this would be our reference channel. The program than measured objects in the other channel(s) that overlapped partially or completely with the 488-positive areas. For the analysis, well-formed protrusions of cells at the wound edge were cropped out as areas to be analyzed. On occasion, there were areas that contained 10 or more objects. These were ROIs with oversaturated pixels that were discarded from the analysis. Mean values, STDEV, and *n* values were then used to calculate P-values in unpaired student’s *t* tests using GraphPad QuickCalcs (GraphPad Software). Graphs were prepared using GraphPad Prism software.

### Electron Microscopy: pre-Embedding Immunolabeling, Silver Enhancement, and Imaging

After 3 washes of the specimens in PBS^2+^ (PBS plus Ca^2+^ and Mg^2+^, see section on immunofluorescence staining), specimens were fixed with 8% PFA (EM grade) in 0.25 M HEPES buffer (pH about 7.1) for 1-h at RT. New fixative was then added followed by an overnight incubation at 4°C. The next day, specimens were washed 3 x for 5 min each in PBS at RT with gentle shaking on a rocker. Specimens were then quenched for 15 min in 50 mM NH_4_Cl in PBS, followed by 3 washes in PBS, 5 min each, at RT with gentle shaking. Specimens were blocked with 2% goat serum, 0.1% BSA-c (from Aurion), and 0.2% saponin in PBS for 1-h at RT. Anti-HA antibodies were diluted (1:75) in block solution and incubated with specimens for 1-h at RT followed by 4.5-h at 4°C. Subsequently, specimens were washed 3 x in PBS, 5 min each, at RT with gentle shaking. 1.4-nm gold-labeled secondary goat anti-mouse antibodies were diluted (1:50) in block solution and incubated with the specimens overnight at 4°C. The next day, specimens were washed 3 x in PBS, 5 min each, at RT with gentle shaking. After the last wash, specimens were incubated with 2% glutaraldehyde in PBS for 30 min at RT, followed by 3 washes in PBS, 5 min each, at RT with gentle shaking. Specimens were quenched for 5 min in 50 mM NH_4_Cl in deionized water at RT, followed by 2 washes in deionized water, 5 min each, at RT.

The subsequent silver enhancement reaction was carried out in a darkroom. The developer was prepared according to the manufacturer’s instruction and 50-µl drops were added to the probes. After 10 min, the reaction was stopped by 6 washes with deionized water, 2 min each, at RT. Samples were immediately post-fixed in 1% osmium tetroxide and 3% uranyl acetate, dehydrated in a series of ethanol washes and embedded in Epon 812 resin. The resin blocks were cured for 1 d at 56°C, and glass coverslips were removed from the blocks in liquid nitrogen. Ultrathin 70-nm sections were cut using a Leica UC7 Ultratome, deposited on copper grids and contrasted with Reynolds lead citrate and uranyl acetate. EM data was gathered with a Hitachi HD-2300A Dual EDS Cryo STEM operated at 200 kV utilizing the phase contrast bright field (TE) detector and the high angle annular dark field (HAADF) detector. Images were collected on Gatan Digital Micrograph with a Digiscan system. Images were processed and combined using Adobe Photoshop and Adobe Illustrator.

### Western Blot Analysis and Co-Immunoprecipitations

For Western blot analysis of total cell lysates, cells were seeded at a (1:1) dilution into 6-well plates and grown for 1 d. Cells were washed 2 x with ice-cold PBS^2+^ (PBS plus Ca^2+^ and Mg^2+^, see section on immunofluorescence staining) and 600 µl RIPA buffer (50 mM Tris-HCl, pH 7.6, 150 mM NaCl, 1% Triton X-100, 0.5% deoxycholate, 0.1% SDS, and 1 x protease inhibitors [Roche]) was added per well. Subsequently, cells were scraped and passaged 3 x through a 22 ½ G needle and 1-ml syringe, followed by a 30-min incubation on ice. Samples were then spun at 16,100 × *g* for 15 min at 4°C (microcentrifuge, Eppendorf). Supernatants were transferred to new tubes, and protein concentrations were determined using the BCA protein assay (Thermo Fisher Scientific). Protein lysates were analyzed by SDS PAGE and Western blot using fluorescently-labeled secondary antibodies. Specimens were imaged using a Li-Cor Odyssey Blot imager. Representative exposures were assembled and combined using Adobe Photoshop and Adobe Illustrator.

For co-immunoprecipitations, cells were seeded (1:1) into 10-cm plates 1 day prior to the experiment. Cells were washed 2 x with PBS^2+^ and 4 ml lysis buffer (50 mM Tris-HCl, pH 7.4, 300 mM NaCl, 1% Triton X-100, 0.1% BSA, 1 x protease inhibitors [Roche]) was added to the washed samples. Cells were scraped with a cell scraper and passed 3 x through a 22 ½ G needle and 5-ml syringe. Lysates were incubated on ice for 30 min, and spun at 16,100 × *g* for 15 min at 4°C (microcentrifuge, Eppendorf). Lysis supernatants were then incubated for 1-h at 4°C, rotating end-over-end, with relevant antibodies that were pre-bound to protein G-Sepharose beads. Co-immunoprecipitates were washed 2 x with lysis buffer, and 1 x with lysis buffer without Triton X-100. Samples were analyzed by SDS PAGE and Western blot using HRP-labeled secondary antibodies and SuperSignal West Pico chemiluminescent substrate (Thermo Fisher Scientific). Films with representative exposure times were scanned. Data was then assembled and combined using Adobe Photoshop and Adobe Illustrator.

For immunoisolations of clathrin-coated vesicles, cells were split (1:1) into six 20-cm dishes. Typically 2 d after seeding, cells were wounded by scratching the monolayers multiple times with a P1000 pipette tip and incubated for 1.5 h. Cells were then washed twice with PBS^2+^, and 2 ml buffer D (10 mM HEPES, 150 mM NaCl, 0.5 mM MgCl_2_, 1 X protease inhibitors [mini tablets from Pierce], and 0.02 [wt/vol] NaN_3_) was added. Cells were scrapped off the plate, combined and homogenized with a cell cracker followed by a clarifying spin in an Allegra D centrifuge (Beckman) at 4°C for 30 min at 5,000 × *g*. Supernatants were harvested and adjusted with Triton X-100 to a final concentration of 1%, and spun for 1 hour in a MAX-E tabletop ultracentrifuge (Beckman) at 4°C and 100,000 × *g* (47,000 rpm, MLA80 rotor). Pellets were resuspended in 3 ml buffer D^+^ (buffer D plus 250 mM sucrose) and subjected to immunoprecipitations with goat anti-GFP antibodies (generic goat IgGs as control) or mouse anti-γ-adaptin antibodies (mouse hybridoma C7 antibodies as control) bound to protein G-Sepharose beads (Fisher Scientific). Samples were rotated end-over-end at 4°C for 2 h and washed 2 x in buffer D^+^ and 1 x in PBS. Immunoprecipitates were denatured in SDS sample buffer, vigorously shaken for 5 min at RT, boiled and analyzed by SDS PAGE and Western blot using fluorescently-labeled secondary antibodies followed by analysis using a Li-Cor Odyssey Blot imager. Representative exposures were assembled and combined using Adobe Photoshop and Adobe Illustrator.

## ACKNOWLEDGEMENTS

We thank Dr. Linton Traub for anti-µ1 antibodies, Samuel Romo for technical assistance, and our Northwestern colleagues Drs. Constadina Arvanitis, David Kirchenbuechler, Farida Korobova, Wensheng Liu, Joshua Rappoport (now at Boston College), Eric Roth, and Lili Zheng for advice and discussions.

This work was supported by an A*STAR Singapore postdoctoral fellowship to S.F. Ang, by the National Institutes of Health grant GM070736 to H. Fölsch, and cores of the Skin Disease Research Center (grant P30AR-57216) and Center for Advanced Microscopy (grants NCI CCSG P30 CA060553 awarded to the Robert H Lurie Comprehensive Cancer Center and NCRR 1S10 RR031680). Further, this work made use of the BioCryo facility of Northwestern University’s NU*ANCE* Center, which has received support from the Soft and Hybrid Nanotechnology Experimental (SHyNE) Resource (NSF ECCS-1542205); the MRSEC program (NSF DMR-1720139) at the Materials Research Center; the International Institute for Nanotechnology (IIN); and the State of Illinois, through the IIN. It also made use of the CryoCluster equipment, which has received support from the MRI program (NSF DMR-1229693).

The authors declare no competing financial interests.

Author contributions: Conceptualization: S.F. Ang, M.J. Kell, and H. Fölsch; formal analysis: S.F. Ang, M.J. Kell, L. Pigati, and H. Fölsch; investigation: S.F. Ang, M.J. Kell, L. Pigati, A. Halpern, and H. Fölsch; writing original draft: H. Fölsch; editing and review: S.F. Ang, M.J. Kell, L. Pigati, A. Halpern, and H. Fölsch; visualization: S.F. Ang, M.J. Kell, and H. Fölsch; supervision: H. Fölsch; funding acquisition: S.F. Ang, and H. Fölsch.

## ABREVIATIONS

AP, adaptor protein; ARH, autosomal recessive hypercholesterolemia protein; CHC, clathrin heavy chain; EM, electron microscopy; GAPDH, glyceraldehyde 3-phosphate dehydrogenase; HBE, human bronchial epithelial; LLC-PK1, Lilly laboratories cell porcine kidney; MDCK, Madin-Darby canine kidney; PI(3,4,5)P_3_, phosphatidylinositol 3,4,5-trisphosphate; qRT-PCR, quantitative real-time reverse transcription PCR; RE, recycling endosome; shRNA, short hairpin RNA; STORM, stochastic optical reconstruction microscopy; TIRF, total internal reflection fluorescence; YFP, yellow fluorescent protein

## VIDEO AND SUPPLEMENTAL FIGURE LEGENDS

**Video S1: Live TIRF imaging of AP-1B-YFP during wound healing**

LLC-PK1::µ1B-YFP cells were grown in matrigel-coated MatTek dishes, wounded and imaged with live-TIRF settings for a total of 30 sec. Green arrows at the start of the movie point to spots present at the beginning but not at 15 sec, and orange arrows in the middle of the movie point to spots present at around 15 sec but not at the beginning of the movie. Signal intensities were equalized to prepare this video. Bar is 10 µm.

**Video S2: Live Biostation imaging of LLC-PK1::µ1A cells**

LLC-PK1::µ1A cells were grown in matrigel-coated MatTek dishes for 2d, wounded and imaged for 4 h in a Biostation. Images were acquired every 15 min. Bar is 50 µm.

**Video S3: Live Biostation imaging of LLC-PK1::µ1B cells**

LLC-PK1::µ1B cells were grown in matrigel-coated MatTek dishes for 2d, wounded and imaged for 4 h in a Biostation. Images were acquired every 15 min. Bar is 50 µm.

**Figure S1:**
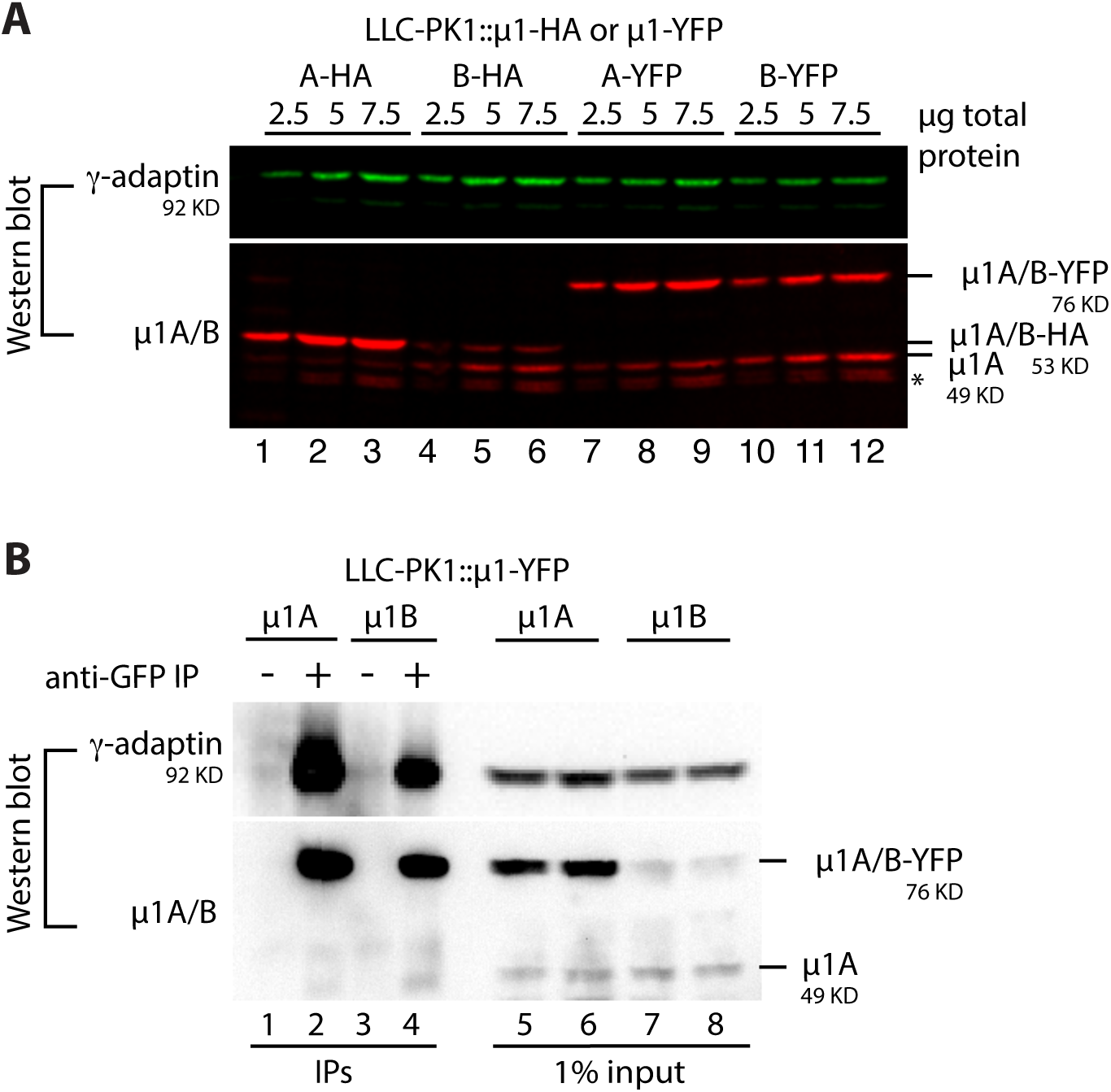
µ**1**A**-YFP and µ1B-YFP are incorporated into AP-1 complexes** (A) Cell lysates of LLC-PK1 cells stably expressing µ1A-HA (A-HA, lanes 1 – 3), µ1B-HA (B-HA, lanes 4 – 6), µ1A-YFP (A-YFP, lanes 7 – 9), and µ1B-YFP (B-YFP, lanes 10 – 12) were analyzed by SDS PAGE and Western blot using primary antibodies specific for *γ*-adaptin and an anti-µ1A/B antibody followed by fluorescently-labeled secondary antibodies. * indicates a degradation band. (B) LLC-PK1::µ1A-YFP (lanes 1 & 2) and LLC-PK1::µ1B-YFP (lanes 3 & 4) cell lysates were subjected to co-immunoprecipitations with antibodies directed against GFP (lanes 2 & 4), or control antibodies (C7 raised against LDL receptor, lanes 1 & 3). Immunoprecipitates were analyzed by Western blot using primary antibodies against *γ*-adaptin and against µ1A/B followed by HRP-labeled secondary antibodies. 1% input for lanes 1 & 2 are shown in lanes 5 & 6, and 1% input for lanes 3 & 4 are shown in lanes 7 & 8.

